# Endothelial cell derived microvesicles modulate cerebrovascular and brain aging via miR-17-5p and serve as potential biomarkers for vascular aging

**DOI:** 10.1101/2023.02.02.526751

**Authors:** Huiting Zhang, Zi Xie, Yuhui Zhao, Yanyu Chen, Xiaobing Xu, Ye Ye, Yuping Yang, Wangtao Zhong, Bin Zhao, Xiaotang Ma

## Abstract

Vascular aging, characterized by brain endothelial cells (ECs) senescence and dysfunction, has been known to lead to various age-related cerebrovascular and neurodegenerative diseases. However, its underlying mechanisms remain elusive. ECs derived microvesicles (EMVs) and exosomes (EEXs) carry the characteristics of parent cells and transfer their contents to modulate the functions of recipient cells, holding the potential to evaluate or regulate vascular aging. Here, we found that young or aged ECs released EMVs were more effective than their released EEXs on alleviating or aggravating mice cerebrovascular and brain aging as indicated by SA-β-gal staining, cerebral blood flow, blood brain barrier function, aging related markers and cognitive ability test. We further identified that these EMVs regulated cerebrovascular and brain aging by transferring miR-17-5p and could modulate ECs senescence and functions via miR-17-5p/PI3K/Akt pathway. Plasma EMVs and their contained miR-17-5p (EMV-miR-17-5p) were significantly increased or decreased in the elderly, and were closely correlated with vascular aging. Receiver Operating Characteristic (ROC) analysis showed that the area under the curve was 0.724 for EMVs, 0.77 for EMV-miR-17-5p and 0.815 for their combination for distinguishing vascular aging. Our results revealed the novel roles for EMVs that could more effectively modulate vascular and brain aging than EEXs by regulating ECs functions through miR-17-5p/PI3K/Akt pathway, and also suggested that EMVs and EMV-miR-17-5p represent promising biomarkers and therapeutic targets for vascular aging.

## Introduction

Aging and aging-related diseases have become the most important medical and social issue worldwide. It is estimated that more than 20% of global population will be over 60 years old by 2050 [1]. Vascular aging is closely associated with the aging process, which could cause insufficient blood flow to organs, resulting in organ atrophy and dysfunction [2]. Vascular aging in the central nervous system can induce brain aging manifested with cognitive and sensorimotor decline and even neurodegenerative diseases [3, 4]. Thus, monitoring and preventing vascular aging is important for aging-related diseases, especially for brain. To date, there still lack of effective approaches to evaluate or alleviate vascular and brain aging.

Vascular aging is manifested by endothelial cells (ECs) senescence and ECs dysfunction in the early stage [1]. During vascular aging, ECs proliferation, migration, oxidative stress and apoptosis are dysregulated [5], along with accumulated senescent ECs [1], resulting in vascular vulnerability and related diseases like atherosclerosis, cerebrovascular disease, and degenerative impairment [1, 6]. Theoretically, ECs are primary targets for reflecting and regulating vascular aging. Extracellular vesicles, which mainly consist of microvesicles and exosomes, are released by various cells in response to activation or apoptosis [7, 8]. These vesicles carry the characteristic of parent cells and elicit diverse effects on recipient cells by transferring their contents [9, 10]. Endothelial microvesicles (EMVs) and endothelial exosomes (EEXs) are important mediators of intercellular communication [11], and also could be novel markers of ECs dysfunction or damage [7, 9]. The release and functions of EMVs/EEXs mainly depend on the stimulation and the status of ECs. Under normal condition, EMVs/EEXs could promote viability and inhibit apoptosis of human brain ECs [12, 13]. Hyperglycemic-induced EMVs or atheroprone low-oscillatory shear stress induced EEXs caused higher levels of apoptosis of human umbilical vein endothelial cells (HUVECs) [14, 15]. The plasma level of EMVs has been shown to significantly elevate in the elderly [16]. The EMVs and EEXs of HUVECs are increased in release when cells undergo senescence, and these EMVs can induce calcification of human aortic smooth muscle cells and apoptosis and senescence of normal ECs [17-19]. An earlier study has found that blood transfusion of young mice reverses age-related impairments in cognitive function and synaptic plasticity in old mice [20], raising the possibility that young EEXs or EMVs might serve as a novel cell-free source to challenge aging. To be noted, the biogenesis of EMVs and EEXs is different, and their effects on the recipient cells/tissues await for validation [7, 9]. Based on these, we hypothesized that EMVs or EEXs might play critical roles in regulating cerebrovascular and brain aging and ECs functions, and hold biomarker potential for vascular aging.

MicroRNAs (miRNAs) are crucial molecules regulating cell senescence and diverse functions and are the main functional cargo of EMVs/EEXs [21]. For example, the level of EMVs containing miR-126 is significantly decreased in patients with diabetes mellitus [22]. MiR-126 enriched EMVs promote the migration and proliferation of human coronary artery ECs [22]. MiR-19a-3p in EEXs derived from HUVECs promoted ECs tube formation and proliferation [23]. Thus, it is of great interest that which miRNA is important in EMVs/EEXs in regulating vascular aging.

In this study, we aimed to investigate the effects and discrepancy of EMVs and EEXs on regulating cerebrovascular and brain aging, and to explore the underlying miRNAs mechanisms and the biomarker potential. First, we determined the effects of EMVs and EEXs derived from young or old mouse brain ECs on cerebrovascular and brain aging. Based on our finding that EMVs were more effective, we further investigated the effects of EMVs on ECs functions in vivo and in vitro, identified miR-17-5p as the key functional miRNA in EMVs and analyzed the related pathways to explore the underlying mechanisms. In clinical study, we measured the levels of plasma EMVs and their contained miRNA in people of different ages for their association with vascular aging.

## Results

### Character of EMVs and EEXs

Replicative aging model with postnatal-derived brain microvascular endothelial cells (BMECs) was established, and BMECs were identified by immunofluorescence. Results showed that BMECs were positive for ECs markers CD31^+^ and the purity was over 90% (**Fig 1A**). As shown in **Fig 1B**, senescence-associated β-galactosidase (SA-β-gal) staining analysis showed that senescent BMECs (o.cells) displayed higher percentage of senescent cells compared with young BMECs (y.cells). EMVs and EEXs purified from conditioned medium collected from y.cells (y.EMVs, y.EEXs) and o.cells (o.EMVs, o.EEXs) were characterized by nanoparticle tracking analysis (NTA) and transmission electron microscopy (TEM) which showed that the sizes of y.EMVs/o.EMVs and y.EEXs/o.EEXs were 200-800 nm and 50-100 nm in diameter, respectively (**Fig 1C and 1D**).

**Fig 1.**
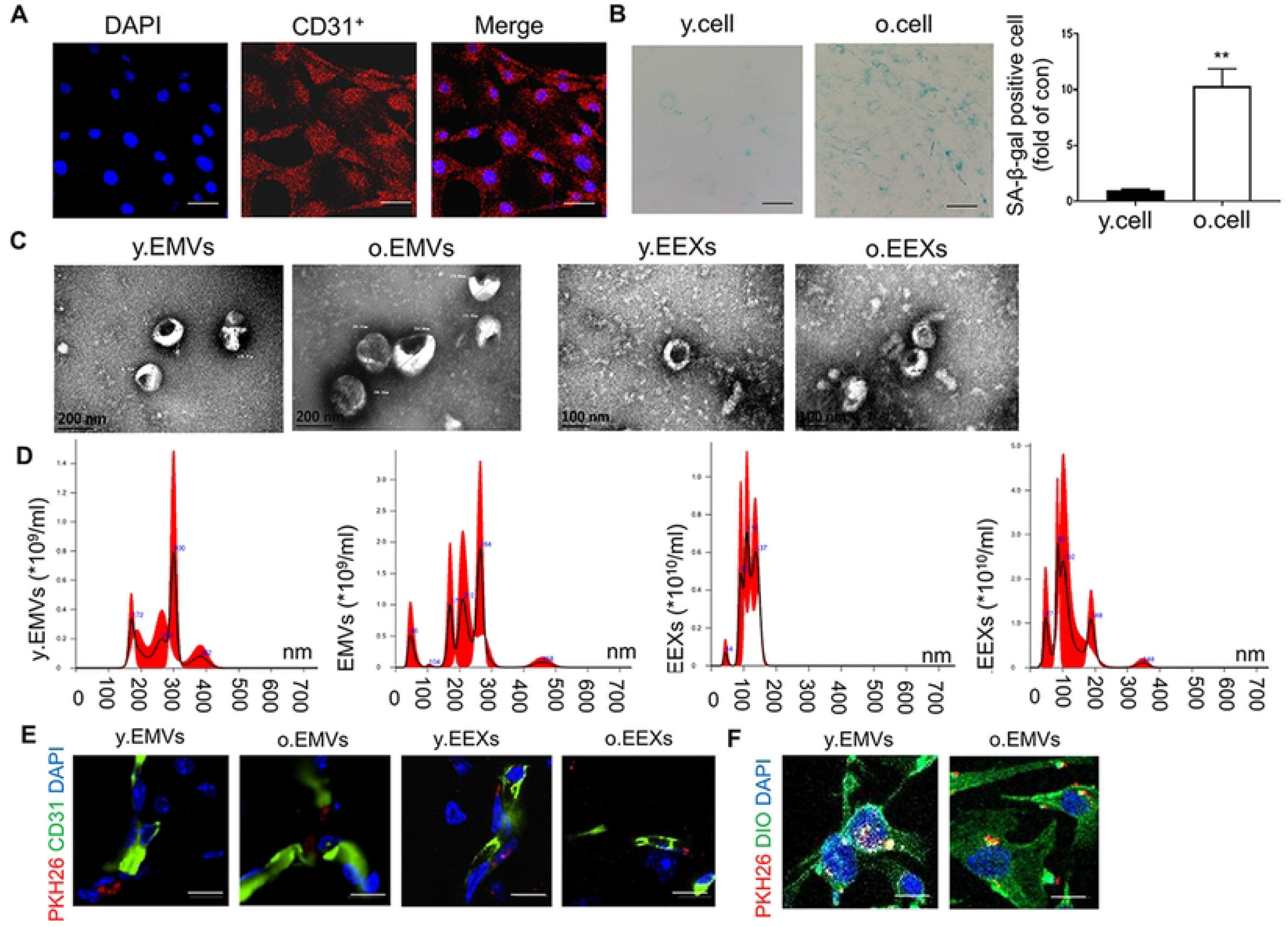
Identification of BMECs and the characterization of EMVs/EEXs from y.cells and o.cells, and the internalization of EMVs/EEXs in BMECs and mouse brain vasculature. (A) Immunofluorescent characterization of BMECs. The isolated BMECs were 95% positive for CD31(red), confirming the epithelial nature of these cells (scale bar=30 μm, *n* = 3 technical replicates). (B) Identification of senescence of BMECs analyzed by SA-β-gal staining (scale bar=200 μm). Senescent BMECs showed higher percentage of SA-β-gal positive stained cells marked blue (>60%) than young BMECs (Student’s *t*-test, mean ± SEM, *n* = 6 technical replicates, ^**^ *p* < 0.01). (C) Representative transmission electron microscopy (TEM) analysis of y.EMVs/o.EMVs and y.EEXs/o.EEXs (scale bar=200 nm for EMVs, scale bar=100 nm for EEXs, *n* = 3 technical replicates). (D) Average size distribution and concentration of y.EMVs/o.EMVs and y.EEXs/o.EEXs determined by nanoparticle tracking assay (*n* = 3 technical replicates). (E) Confocal microscopy images of the internalization of EMVs and EEXs labeled with PKH 26 (marked red) in mouse brain vasculature, nuclei were stained with DAPI (marked blue), blood vessels were stained with CD31^+^ (marked green), scale bar=20 μm, *n* = 6 biological replicates per group. (F) Confocal microscopy images of the internalization of EMVs labeled with PKH 26 (marked red) in BMECs, nuclei were stained with DAPI (marked blue), cell membrane was stained with DIO (marked green), scale bar=20 μm. *n* = 3 technical replicates.

### Y.EMVs mitigated while o.EMVs aggravated cerebrovascular and brain aging more significantly than y.EEXs or o.EEXs respectively

After verified that EMVs and EEXs could be internalized into mouse brain vasculature (**Fig 1E**), we measured their effects on cerebrovascular aging. SA-β-gal staining in basilar artery was utilized to evaluate cerebrovascular aging. Results showed that the number of senescent cells in basilar artery was dramatically 4.5-fold higher in o.mice than that in y.mice (**Fig 2A, 2B and 2G**). Y.EMVs and y.EEXs significantly decreased 60% and 45% the senescent cell number of basilar artery in o.mice (**Fig 2B and 2G**). O.EMVs and o.EEXs significantly increased senescent cell number of basilar artery in y.mice (**Fig 2A and 2G**), the effect of y.EMVs/o.EMVs was more significant than y.EEXs/o.EEXs (**Fig 2G**). Cerebral blood flow (CBF) and blood brain barrier (BBB) have been reported as indicators of cerebrovascular aging [4]. We found that CBF was dramatically decreased about 50% in o.mice compared with y.mice (**Fig 2C, 2D and 2H**). Y.EEXs and y.EMVs significantly increased CBF as evidenced by a 1.46 and 1.88-fold elevation, and the increase of y.EMVs was more significant (**Fig 2D and 2H**). The results were further supported by 2.8 and 1.1-fold increase of cerebral microvascular density (cMVD) in y.EMVs and y.EEXs groups, respectively (**Fig 2F and 2I**). In contrast, o.EMVs and o.EEXs reduced CBF of y.mice by 65% and 55% respectively (**Fig 2C and 2H**), and o.EMVs was more significant in decreasing cMVD of y.mice (61%) than o.EEXs (44%) (**Fig 2E and 2I**). Evans Blue (EB) extravasation was applied to detect BBB function. Results showed that EB leakage was 1.6 times higher in o.mice compared with y.mice. Y.EMVs and y.EEXs significantly improved while o.EMVs and o.EEXs insulted BBB integrity, the effects of y.EMVs/o.EMVs were more significant (**Fig 2J**).

**Fig 2.**
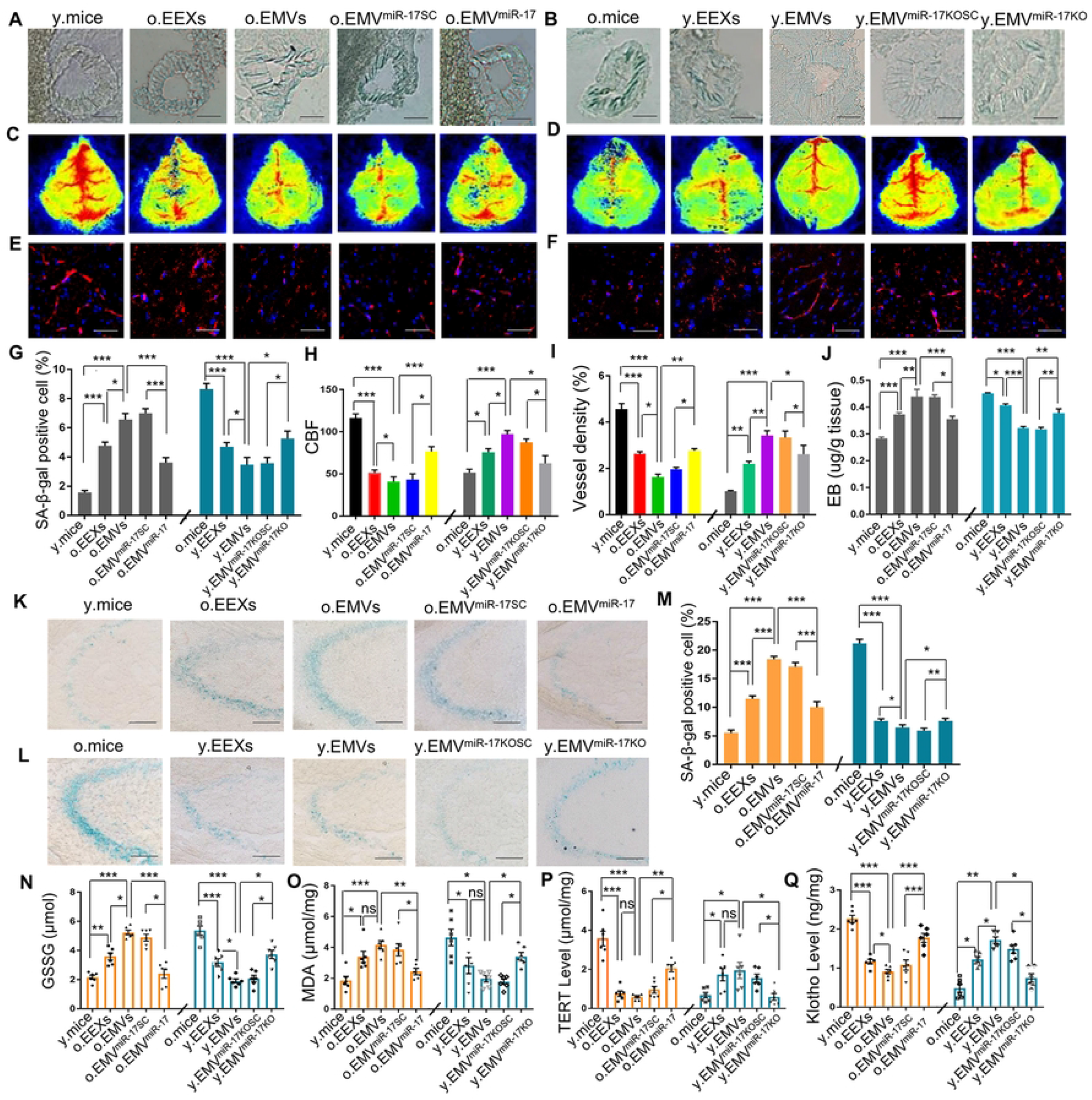
Y.EMVs mitigated while o.EMVs aggravated cerebrovascular and brain aging more significantly than y.EEXs or o.EEXs by modulating miR-17-5p. (A, B) Representative images of SA-β-gal staining in basilar artery (Scale bar=20 μm, *n* = 6 biological replicates per group). (C, D) Representative images of CBF tested by laser doppler flowmetry in diverse groups (*n* = 10 biological replicates per group). (E, F) Confocal microscopy images of cerebral microvascular density marked with CD31^+^ in mice brain. Red, CD31^+^; blue, DAPI for nuclei staining (scale bar=50 μm, *n* = 6 biological replicates per group). (G) The percentage of SA-β-gal staining positive cell in basilar artery (one-way ANOVA, Tukey’s multiple comparisons test, mean ± SEM, *n* =6 biological replicates per group, ^*^ *p* < 0.05, ^***^ *p* < 0.001). (H) Quantification of CBF in various groups (Kruskal–Wallis ANOVA, post hoc Dunn’s multiple comparison test, mean ± SEM, *n* = 10 biological replicates per group, ^*^ *p* < 0.05, ^***^ *p* < 0.001). (I) Quantification of vessel density in various groups (one-way ANOVA, Tukey’s multiple comparisons test, mean ±SEM, *n* = 6 biological replicates per group, ^*^ *p* < 0.05, ^**^ *p* < 0.01, ^***^ *p* < 0.001). (J) EB extravasation in mice brains (one-way ANOVA, Tukey’s multiple comparisons test, mean ± SEM, *n* = 4 biological replicates per group, experiment repeated for 3 times, ^*^ *p* < 0.05, ^**^ *p* < 0.01, ^***^ *p* < 0.001). (K, L) Representative images of SA-β-gal positive cells in CA3 region of hippocampus in diverse groups (scale bar=100 μm. *n* = 6 biological replicates per group). (M) Quantification of SA-β-gal positive cell in various groups (one-way ANOVA, Tukey’s multiple comparisons test, mean ± SEM, *n* = 6 biological replicates per group, ^*^ *p* < 0.05, ^**^ *p* < 0.01, ^***^ *p* < 0.001). (N) Plasma GSSG expression of mice in diverse groups (one-way ANOVA, Tukey’s multiple comparisons test, mean ± SEM, *n* = 6 biological replicates per group, ^*^ *p* < 0.05, ^**^ *p* < 0.01, ^***^ *p* < 0.001). (O-Q) MDA, TERT and klotho expression in mice brain (one-way ANOVA, Tukey’s multiple comparisons test, mean ± SEM, *n* = 6 biological replicates per group, ^*^ *p* < 0.05, ^**^ *p* < 0.01, ^***^ *p* < 0.001, *ns*, no significance).

As cerebrovascular aging contributes to brain aging [24], we detected the effects of EMVs and EEXs on brain aging. First, we confirmed that the senescent cell number in hippocampus CA3 region of o.mice was dramatically increased (**Fig 2K-2M**). O.EMVs and o.EEXs increased about 3.3-fold and 1.1-fold of SA-β-gal positive cells in the CA3 region of y.mice, and o.EMVs was more significant (**Fig 2K and 2M**). The senescent cell number in o.mice treated with y.EMVs and y.EEXs was decreased by about 72% and 60%, respectively (**Fig 2L and 2M**). The age-related biomarkers oxidized glutathione (GSSG) and malondialdehyde (MDA) were significantly increased, while telomerase reverse transcriptase (TERT) and klotho were decreased in o.mice (**Fig 2N-2Q**). O.EMVs and o.EEXs evidently increased plasma GSSG and brain MDA levels (**Fig 2N and 2O**), while decreased brain klotho and TERT levels in y.mice (**Fig 2P and 2Q**). On the contrary, y.EMVs and y.EEXs played the opposite roles in regulating age-related biomarkers (**Fig 2N-2Q**). The effects of y.EMVs/o.EMVs were more significant than y.EEXs/o.EEXs on regulating the levels of GSSG and klotho.

In furtherance, Morris water maze task was performed to measure the cognitive ability of the mice. We found that o.EMVs and o.EEXs obviously increased the escape latency, decreased time spend in the target quadrant, impaired platform crossing performance and searching strategy of y.mice (**Fig 3A and 3E-3G**). The effects of o.EMVs were more significant than o.EEXs regarding to the escape latency performance (**Fig 3A**). The comparison of o.EMVs and o.EEXs on quadrant time, platform crossing and searching strategy showed no statistical difference (**Fig 3A and 3E-3G**). Y.EMVs dramatically shortened the escape latency (**Fig 3C**), increased quadrant time (**Fig 3E**), improved platform crossing performance and promoted the searching strategy of o.mice (**Fig 3F and 3H**). Y.EEXs significantly promoted platform crossing performance (**Fig 3F**) and searching strategy (**Fig 3H**), but barely influenced the escape latency and quadrant time of o.mice (**Fig 3C and 3E**). The effects of y.EMVs were more evident than y.EEXs on escape latency and searching strategy (**Fig 3C and 3H**). Taken together, these results confirmed that o.EMVs/o.EEXs deteriorated while y.EMVs/y.EEXs attenuated brain aging, and the effects of EMVs were more remarkable than EEXs.

**Fig 3.**
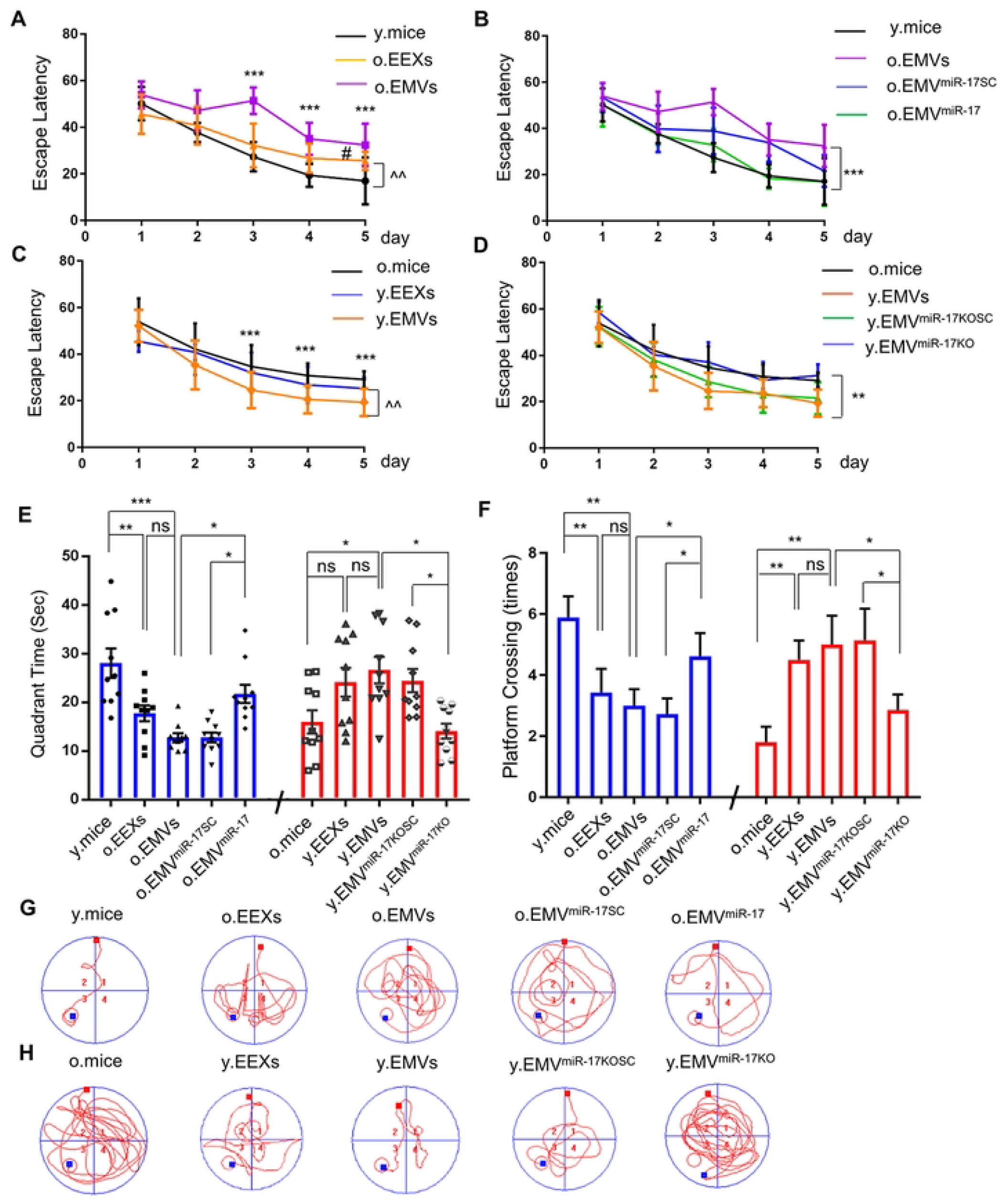
Y.EMVs improved while o.EMVs exacerbated mice cognitive ability more evidently than y.EEXs or o.EEXs by modulating miR-17-5p. (A-D) Escape latency of various groups (two-way ANOVA with repeated measures, mean ± SEM, *n* = 10 biological replicates per group). (E) Time spent in the target quadrant of various groups (one-way ANOVA, Tukey’s multiple comparisons test, mean ± SEM, *n* = 10 biological replicates per group, ^*^ *p* < 0.05, ^**^ *p* < 0.01, ^***^ *p* < 0.001, *ns*, no significance). (F) The numbers of platform crossings of different groups (Kruskal–Wallis ANOVA, post hoc Dunn’s multiple comparison test, mean ± SEM, *n* = 10 biological replicates per group, ^*^ *p* < 0.05, ^**^ *p* < 0.01, *ns*, no significance). (G, H) Searching strategy of diverse groups (*n* = 10 biological replicates per group).

### MiR-17-5p was differentially expressed in y.EMVs and o.EMVs

Considering the effects of EMVs were more significant than EEXs on cerebrovascular and brain aging, we focused on studying the mechanisms of EMVs. MicroRNA sequencing was applied to identify the functional miRNAs in EMVs. Heat map showed 26 significantly downregulated miRNAs and 63 upregulated miRNAs in o.EMVs compared with y.EMVs, and the top ten overexpressed and downregulated genes was presented in the **Fig 4A**. MiR-17-5p was one of the top miRNAs riched in y.EMVs, and previous research has demonstrated its potential role in alleviating aging [25]. Q-PCR results confirmed that the level of miR-17-5p encapsulated in o.EMVs was robustly decreased about 52% compared with y.EMVs (**Fig 4C**). We also found that miR-17-5p levels in o.cells and brain of o.mice were significantly decreased than those in y.cells and brain of y.mice. Y.EMVs dramatically increased miR-17-5p levels in o.cells and brain of o.mice (**Fig 4D and 4E**). On the contrary, o.EMVs decreased miR-17-5p levels in y.cells and brain of y.mice, which is unexpected, and we speculated that o.EMVs might induce a senescent environment to inhibit miR-17-5p expression in recipient cells (**Fig 4D and 4E**). These results implied the role of EMVs might function by regulating miR-17-5p.

**Fig 4.**
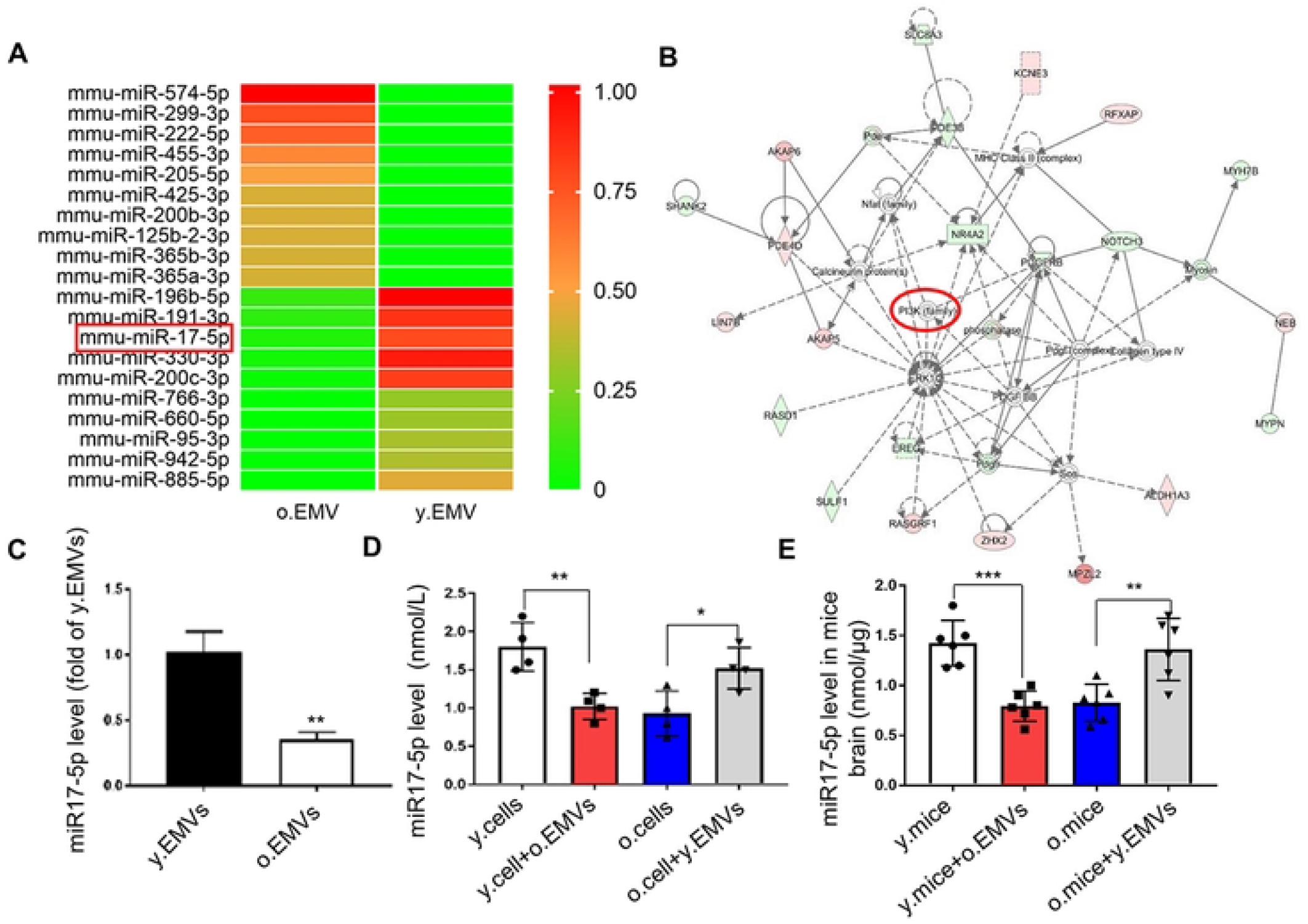
MiR-17-5p was significantly decreased in o.cells, o.mice and o.EMVs and capable of targeting PI3K family. (A) Heat map depicting normalized log_2_ expression values of top 10 up-regulated and 10 down-regulated deferentially expressed genes. Red and green represent high and low relative expression, respectively. (B) MiR-17-5p regulatory network by Ingenuity Pathway Analysis (IPA). (C) MiR-17-5p expression in y.EMVs and o.EMVs (Student’s *t*-test, mean ± SEM, *n* = 5 technical replicates, ^**^ *p* < 0.01). (D) MiR-17-5p expression in different groups (Student’s *t*-test, mean ± SEM, *n* = 4 technical replicates, ^*^ *p* < 0.05, ^**^ *p* < 0.01). (E) Brain miR-17-5p level in diverse groups (Student’s *t*-test, mean ± SEM, *n* = 6 biological replicates per group, ^**^ *p* < 0.01, ^***^ *p* < 0.001).

### Y.EMVs inhibited while o.EMVs induced cerebrovascular and brain aging by modulating miR-17-5p

To clarify the role of miR-17-5p in EMVs, we knocked down miR-17-5p in y.EMVs (y.EMV^miR-17KO^) and overexpressed it in o.EMVs (o.EMV^miR-17^) by infecting y.cells with LV-si-miR-17-5p (y.cells^miR-17KO^) and infecting o.cells with LV-miR-17-5p (o.cells^miR-17^). Results showed that the levels of miR-17-5p in o.cells^miR-17^ and o.EMV^miR-17^ were notably increased (**S1C and S1E Fig**), and the levels of miR-17-5p in y.cells^miR-17KO^ and y.EMV^miR-17KO^ were significantly decreased, as compared with the cells transfected with scrambled controls (**S1D and S1F Fig**).

The results of functional tests showed that o.EMV^miR-17^ had less effect on increasing cell senescent of basilar artery (**Fig 2A and 2G**) and BBB permeability (**Fig 2J**), and on decreasing CBF (**Fig 2C and 2H**) and cMVD (**Fig 2E and 2I**), when compared with o.EMVs and o.EMV^miR-17SC^. Y.EMV^miR-17KO^ decreased senescent number in basilar artery (**Fig 2B and 2G**) and BBB permeability (**Fig 2J**), and increased CBF (**Fig 2D and 2H**) and cMVD (**Fig 2F and 2I**) less effectively than y.EMVs and y.EMV^miR-^ _17KOSC_.

In consistent with their effects on cerebrovascular aging, o.EMV^miR-17^ had weaker impact on upregulating hippocampus senescent cell number (**Fig 2K and 2M**) and the levels of plasma GSSG and brain MDA, and on downregulating levels of brain klotho and TERT compared with o.EMVs and o.EMV^miR-17SC^ (**Fig 2N-2Q**). Moreover, o.EMV^miR-17^ showed less influence on increasing the escape latency (**Fig 3B**), decreasing time spent in the target quadrant and platform crossing times, and deteriorating searching strategy compared with o.EMVs and o.EMV^miR-17SC^ (**Fig 3E-3G**). Y.EMV^miR-17KO^ had less impact on decreasing senescent number and levels of MDA and GSSG, and on increasing TERT and klotho levels compared with y.EMVs and y.EMV^miR-17KOSC^ (**Fig 2L-2Q**). Y.EMV^miR-17KO^ also showed less effect on shortening escape latency, increasing platform crossing times and the time spent in the target quadrant, and improving searching strategy compared with y.EMVs and y.EMV^miR-17KOSC^ (**Fig 3D-3F and 3H**). In summary, these results suggested that EMVs might affect cerebrovascular and brain aging through miR-17-5p modulation.

### Y.EMVs ameliorated while o.EMVs aggravated oxidative stress and apoptosis of mouse brain vasculature via regulating miR-17-5p

As shown in **Fig 5A-5I**, o.EMVs significantly increased the levels of ROS (**Fig 5A and 5C**) and apoptosis (**Fig 5D and 5F**), while decreased NO levels in vasculature of y.mice hippocampal brain slices (**Fig 5G and 5I**). Y.EMVs dramatically decreased the levels of ROS (**Fig 5B and 5C**) and apoptosis (**Fig 5E and 5F**), while enhanced NO generation in vasculature of o.mice hippocampal brain slices (**Fig 5H and 5I**). MiR-17-5p overexpression or downregulation significantly impaired the effects of o.EMVs or y.EMVs and the corresponding scrambled controls (**Fig 5A-5I**).

**Fig 5.**
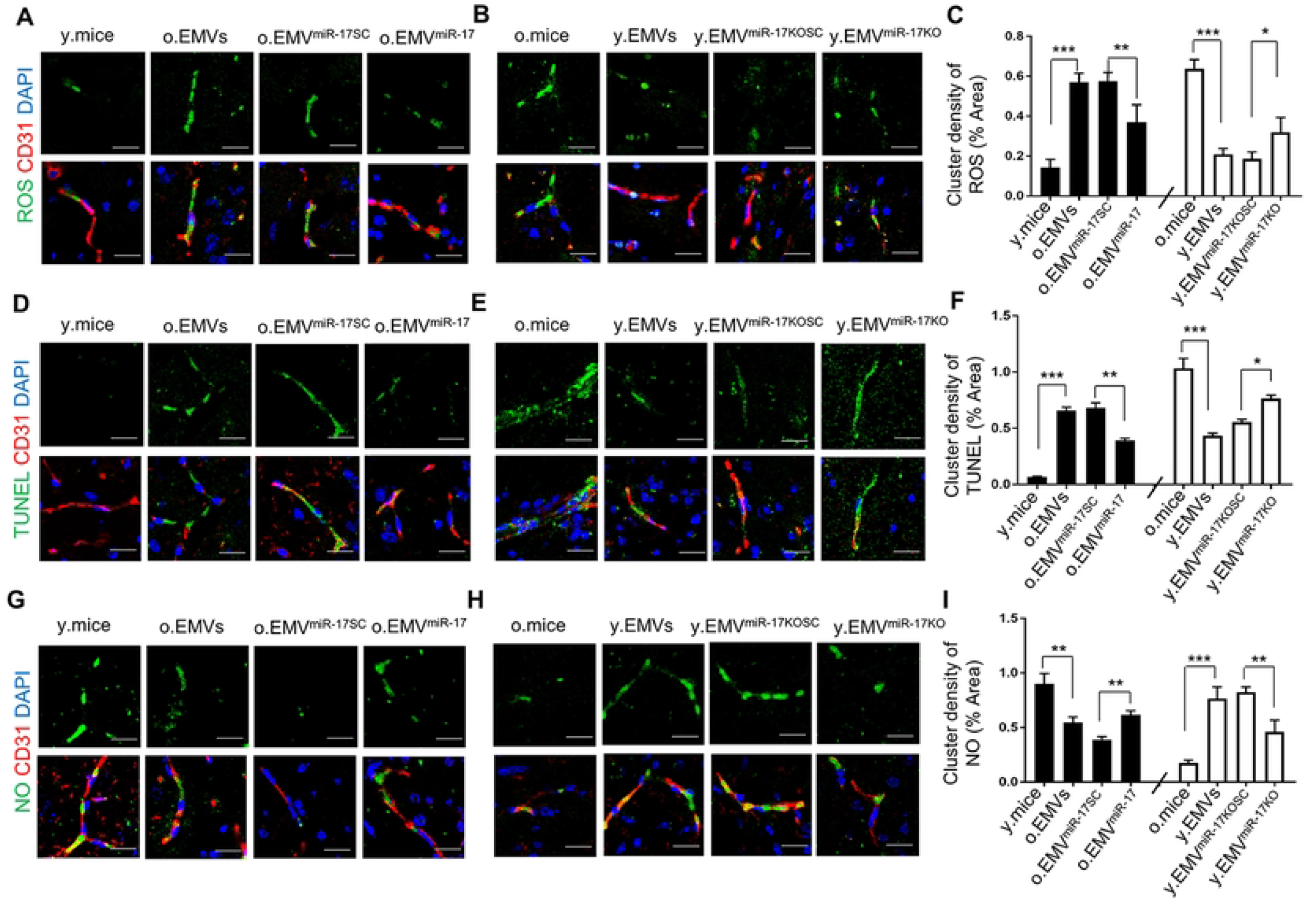
Y.EMVs ameliorated while o.EMVs aggravated oxidative stress and apoptosis in mouse brain vasculature via regulating miR-17-5p. (A, B) Representative images of co-staining of CD31^+^ (marked red) with ROS (marked green) in various groups. CD31^+^ is accepted as endothelium marker, DAPI (marked blue) for nuclei staining (scale bar=30 μm. *n* = 6 biological replicates per group). (C) Summary of ROS expression in mouse brain vasculature (one-way ANOVA, Tukey’s multiple comparisons test, mean ±SEM, *n* = 6 biological replicates per group, ^*^ *p* < 0.05, ^**^ *p* < 0.01, ^***^ *p* < 0.001). (D, E) Immunofluorescent co-staining of CD31^+^ (marked red) with TUNEL (marked green) in mouse brain vasculature in various groups. TUNEL is regarded as apoptosis marker (scale bar=30 μm. *n* =6 biological replicates per group). (F) Summary of apoptotic levels in mouse brain vasculature (one-way ANOVA, Tukey’s multiple comparisons test, mean ± SEM, *n* = 6 biological replicates per group, ^*^ *p* < 0.05, ^**^ *p* < 0.01, ^***^ *p* < 0.001). (G, H) Immunofluorescent co-staining of CD31^+^ (marked red) with NO (marked green) in various groups (scale bar=30 μm. *n* = 6 biological replicates per group). (I) Summary of NO expression in mouse brain vasculature (one-way ANOVA, Tukey’s multiple comparisons test, mean ± SEM, *n* = 6 biological replicates per group, ^**^ *p* < 0.01, ^***^ *p* < 0.001). Data information: For (A-I), same threshold was applied over all pictures.

### Y.EMVs protected while o.EMVs deteriorated BMECs senescence and functions via regulating miR-17-5p/PI3K/Akt pathway

After verifying that EMVs could be incorporated into BMECs (**Fig 1F**), we aimed at investigating whether EMVs exert their effects by regulating BMECs senescence and functions. Results showed that o.EMVs dramatically promoted cell senescence (**Fig 6A and 6C**) and the levels of ROS and apoptosis (**Fig 6D, 6F and 6G**), while decreased NO level (**Fig 6D and 6H**) and the proliferation (**Fig 7D**), migration (**Fig 7A and 7C**) and tube formation ability of y.cells (**Fig 7E and 7G**). O.EMV^miR-17^ showed less deteriorative effects on y.cells compared with o.EMVs and o.EMV^miR-17SC^ (**Figs 6A, 6C, 6D, 6F-6H and 7A, 7C-7E, 7G**). Y.EMVs significantly inhibited cell senescence (**Fig 6B and 6C**), ROS and apoptosis levels (**Fig 6E-6G**), while promoted NO generation (**Fig 6E and 6H**), proliferation (**Fig 7D**), migration (**Fig 7B and 7C**) and tube formation of o.cells (**Fig 7F and 7G**). Y.EMV^miR-17KO^ was less effective than y.EMVs and y.EMV^miR-17KOSC^ (**Figs 6B, 6C, 6E-6H and 7B-7D, 7F, 7G**).

**Fig 6.**
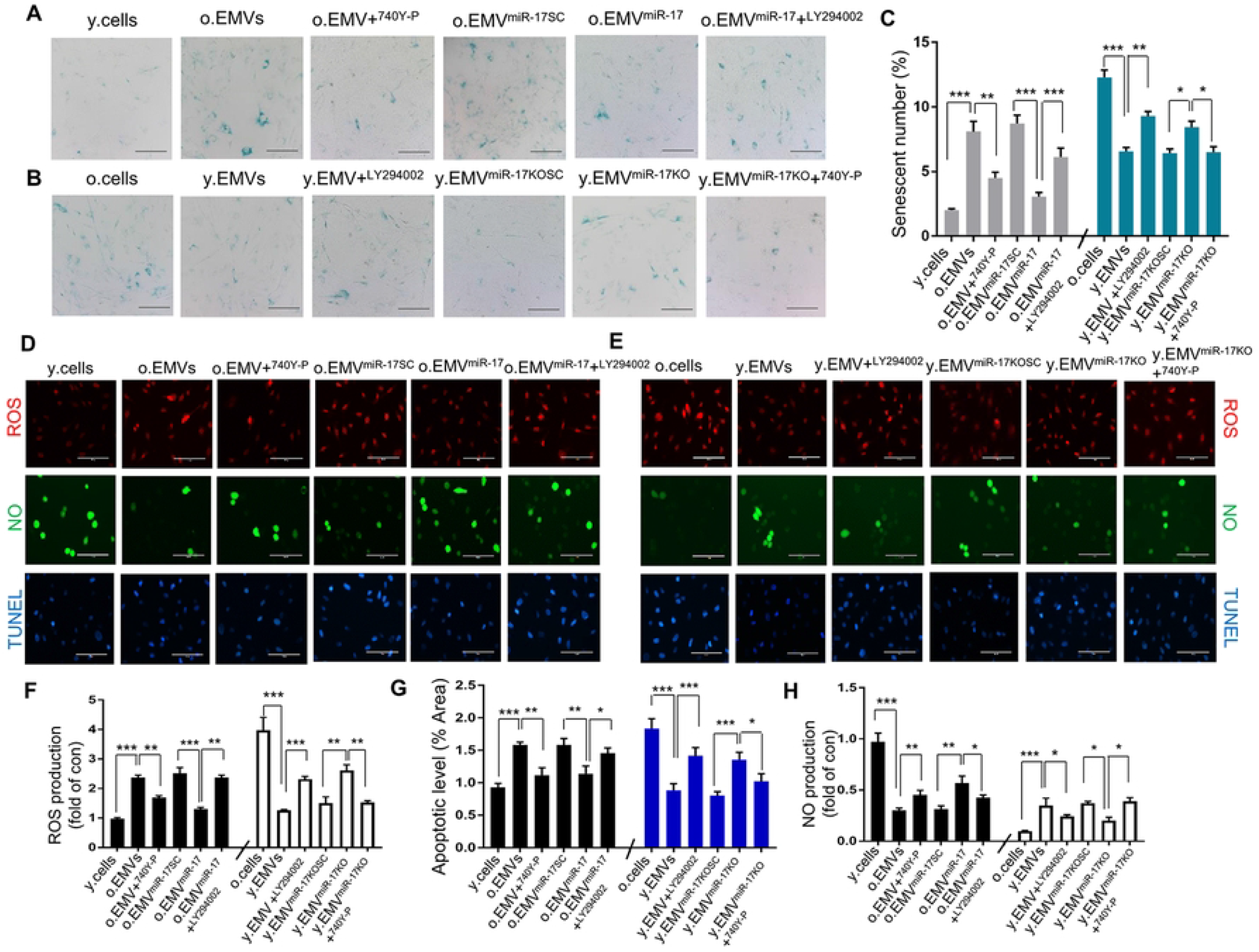
Y.EMVs and o.EMVs modulated BMECs senescence, oxidative stress and apoptosis via the miR-17-5p/PI3K/Akt pathway. (A, B) Representative images of SA-β-gal staining in diverse groups (scale bar=200 μm). (C) Quantification of SA-β-gal staining positive cell in various groups (one-way ANOVA, Tukey’s multiple comparisons test, mean ± SEM, *n* = 5 technical replicates, ^*^ *p* < 0.05, ^**^ *p* < 0.01, ^***^ *p* < 0.001). (D, E) Representative figures of ROS, NO and apoptosis productions in y.cells and o.cells assessed by immunofluorescent dihydroethidium (DHE), DAF-FM DA and TUNEL staining in various groups (scale bar=200 μm). (F-H) Summary of ROS, apoptosis and NO expression in various groups (one-way ANOVA, Tukey’s multiple comparisons test, mean ± SEM, *n* = 5 technical replicates, ^*^ *p* < 0.05, ^**^ *p* < 0.01, ^***^ *p* < 0.001).

**Fig 7.**
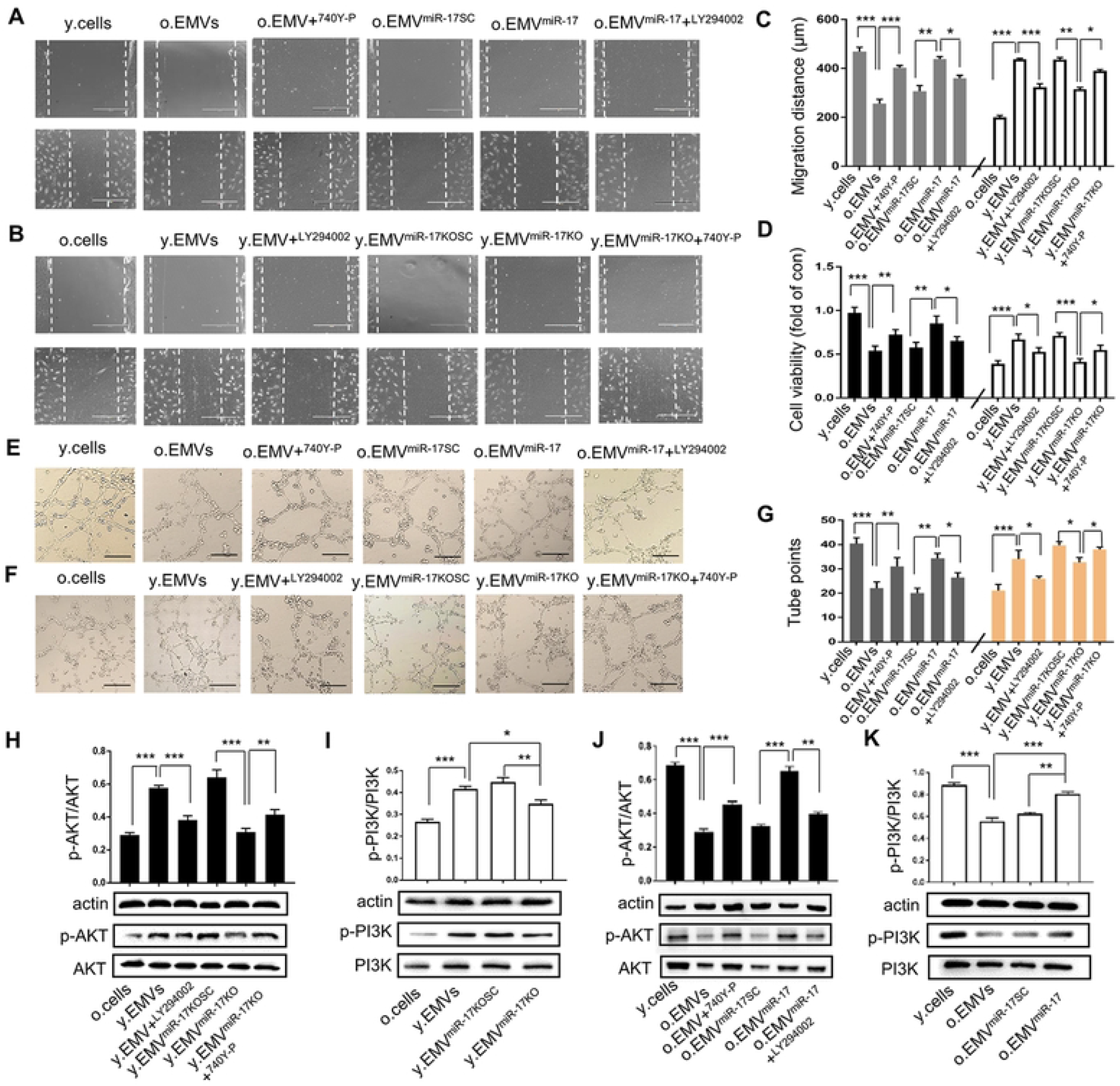
Y.EMVs alleviated while o.EMVs exacerbated BMECs migration, proliferation and tube formation via regulating miR-17-5p/PI3K/Akt pathway. (A, B) Representative images of scratch assay of BMECs migrations (scale bar, 400 μm). (C) Quantification of BMECs migration (one-way ANOVA, Tukey’s multiple comparisons test, mean ± SEM, *n* = 5 technical replicates, ^*^ *p* < 0.05, ^**^ *p* < 0.01, ^***^ *p* < 0.001). (D) CCK-8 assay of BMECs proliferation (one-way ANOVA, Tukey’s multiple comparisons test, mean ± SEM, *n* = 5 technical replicates, ^*^ *p* < 0.05, ^**^ *p* < 0.01, ^***^ *p* < 0.001). (E, F) Representative images of BMECs tube formation (scale bar, 200 μm). (G) Summarized data of tube points per field (one-way ANOVA, Tukey’s multiple comparisons test, mean ± SEM, *n* = 5 technical replicates, ^*^ *p* < 0.05, ^**^ *p* < 0.01, ^***^ *p* < 0.001). (H-K) Western blot was used to analyze the protein levels of p-PI3K/PI3K and p-Akt/Akt in various groups (one-way ANOVA, Tukey’s multiple comparisons test, mean ± SEM, *n* = 5 technical replicates, ^*^ *p* < 0.05, ^**^ *p* < 0.01, ^***^ *p* < 0.001).

As for mechanism study, Ingenuity Pathway Analysis (IPA) results showed that PI3K and its downstream mediator Akt were closely related to miR-17-5p function (**Fig 4B and S1 Table**). Western blot data demonstrated that o.EMVs significantly decreased the levels of p-PI3K/PI3K and downstream p-Akt/Akt in y.cells (**Fig 7J and 7K**). O.EMV^miR-17^ upregulated the levels of p-PI3K/PI3K and p-Akt/Akt compared with o.EMVs and o.EMV^miR-17SC^ (**Fig 7J and 7K**). Y.EMVs dramatically enhanced the levels of p-PI3K/PI3K and p-Akt/Akt in o.cells (**Fig 7H and 7I**). Y.EMV^miR-17KO^ decreased the levels of p-PI3K/PI3K and p-Akt/Akt compared with y.EMVs and the corresponding control (**Fig 7H and 7I**). To further confirm the crucial role of PI3K/Akt pathway in the effects of EMVs and miR-17-5p, the PI3K inhibitor LY-294002 and activator 740Y-P were applied for testing various BMECs functions. 740Y-P significantly ameliorated the impaired effects of o.EMVs on BMECs senescence (**Fig 6A and 6C**) and above mentioned functions (**Figs 6D-6H and 7A-7G**), accompanied with the upregulation of p-Akt/Akt level in BMECs (**Fig 7J**). LY-294002 markedly inhibited the favorable effects of y.EMVs on BMECs senescence (**Fig 6B and 6C**) and functions (**Figs 6D-6H and 7A-7G**), along with the downregulation of p-Akt/Akt levels of BMECs (**Fig 7H**). The effects of o.EMV^miR-17^ treated with LY-294002 on aggravating BMECs senescence (**Fig 6A and 6C**) and functional impairments were more obvious than o.EMV^miR-17^ while less effective than o.EMVs (**Figs 6D-6H and 7A-7G**), accompanied with downregulation of p-Akt/Akt level in BMECs compared with o.EMV^miR-17^ (**Fig 7J**). The effects of y.EMV^miR-17KO^ treated with 740Y-P on inhibiting BMECs senescence (**Fig 6B and 6C**) and functional impairment were marked than y.EMV^miR-17KO^ while weaker than y.EMVs (**Figs 6D-6H and 7A-7G)**, along with upregulation of p-Akt/Akt level in BMECs compared with y.EMV^miR-17KO^ (**Fig 7H**). These results suggested that EMVs regulate BMECs senescence, oxidative stress, apoptosis and angiogenesis by modulating miR-17-5p/PI3K/Akt pathway.

### Clinical subject characteristics

Based on the findings of the differentially expressed EMVs and EMV-miR-17-5p in y.cells/o.cells and y.mice/o.mice, we then investigated the biomarker potential of EMVs and EMV-miR-17-5p in human blood samples. A total of 119 subjects (50 females and 69 males) were enrolled in the study. The subjects were divided into young group (18-45 years old, *n* = 58), middle-age group (46–65 years old, *n* = 33), aged group (> 65 years old, *n* = 28). There were significant differences in age, intima-media thickness (IMT), EMVs and miR-17-5p encapsulated in EMVs (EMV-miR-17-5p) levels among young, middle-age and aged participants (**S2 Table**).

### Plasma levels of EMVs and EMV-miR-17-5p were significantly increased or declined in aged participants

The size of plasma EMVs (defined as CD105^+^CD144^+^ MVs) from young, middle-age and aged people showed no statistical significance (**S2A-S2C Fig**). The level of EMVs in the aged people was approximately 2.5-fold and 1.6-fold higher than those in the young and middle-age groups, and there was no significant difference between young and middle-age groups (**Fig 8A**). These results were in accordance with the data in cultured cells and mice (**S1A and S1B Fig**). The plasma level of EMV-miR-17-5p in the aged group was significantly lower than the young group, but showed no statistical difference compared with middle-age group. The comparison of the levels of EMV-miR-17-5p between the young group and the middle-age group was not statistically significant (**Fig 8B**).

**Fig 8.**
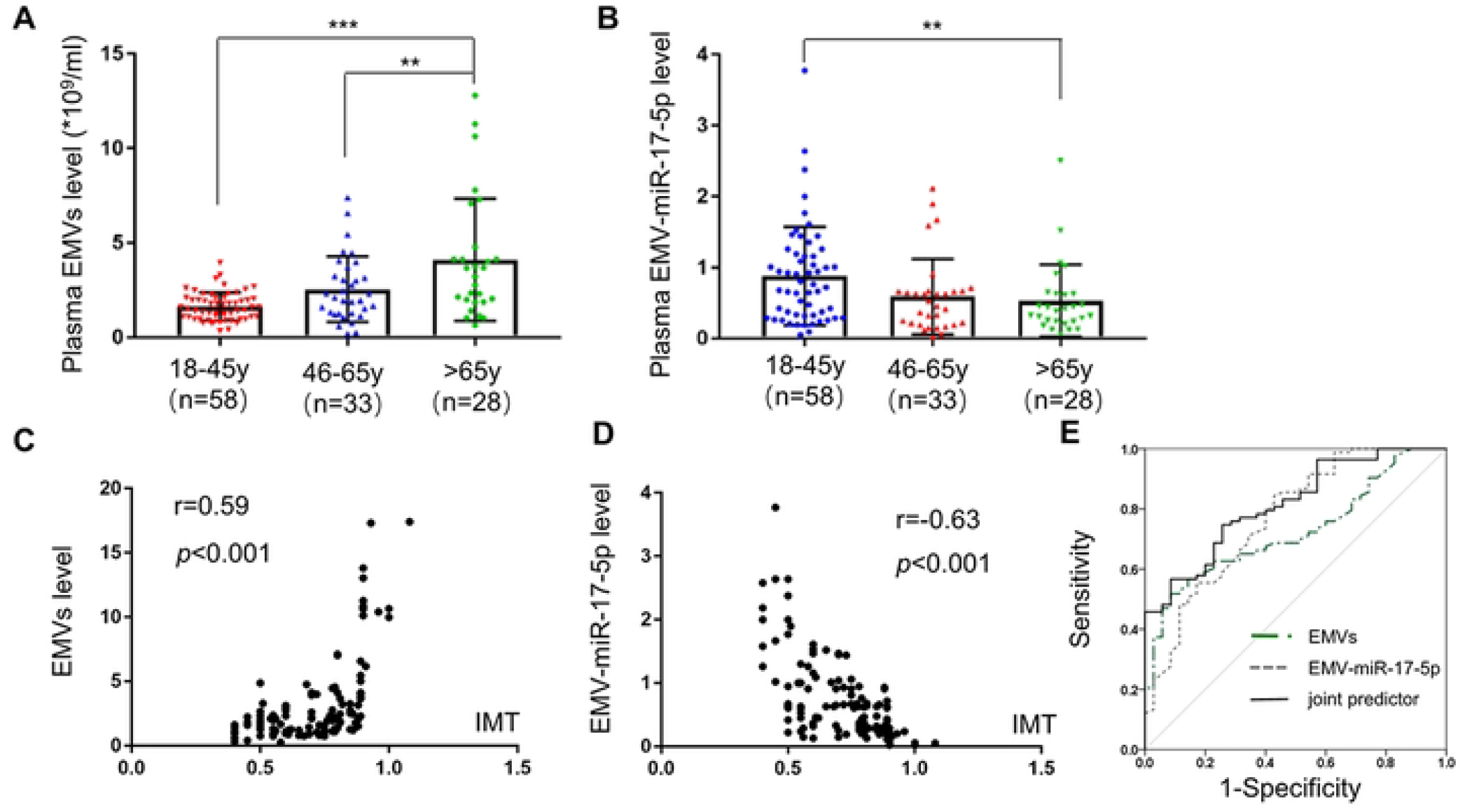
Plasma levels of EMVs and EMV-miR-17-5p were significantly increased or declined with age and were both correlated with vascular aging, and both showed significant diagnostic value for vascular aging. (A) Plasma levels of EMVs in diverse groups tested by NTA (18-45 years old, *n* = 58; 46-65 years old, *n* = 33; >65 years old, *n* = 28) (Kruskal–Wallis ANOVA, post hoc Dunn’s multiple comparison test, mean ± SEM, ^**^ *p* < 0.01). (B) Plasma levels of EMV-miR-17-5p in various groups assessed by q-PCR (Kruskal–Wallis ANOVA, post hoc Dunn’s multiple comparison test, mean ± SEM, ^**^ *p* < 0.01). (C, D) Correlation analysis of levels of EMVs (*n* =119) and EMV-miR-17-5p (*n* = 119) with IMT. Correlations between variables were analyzed by Spearman correlation analysis. (E) ROC analysis was used to assess the diagnostic value of EMVs, EMV-miR-17-5p and their combination for vascular aging defined by IMT. The AUC of EMVs and EMV-miR-17-5p were 0.724 and 0.77, respectively. The AUC of EMVs combined with EMV-miR-17-5p was 0.815.

### Plasma levels of EMVs and EMV-miR-17-5p were correlated with IMT and showed significant diagnostic value for vascular aging

IMT has been considered as an appropriate tool for evaluating vascular aging [1, 26-28]. Spearman correlation analysis revealed a positive moderate correlation between EMVs level and IMT (**Fig 8C**, *r* = 0.59, *P* < 0.001). The level of EMV-miR-17-5p was negatively correlated with IMT (**Fig 8D**, *r* = -0.63, *P* < 0.001).

As shown in **Fig 8E** and **S3 Table**, ROC analysis was used to assess the diagnostic value of EMVs and EMV-miR-17-5p for vascular aging defined by IMT > 0.58 mm [1, 26-28]. The area under the ROC (AUC) of EMVs and EMV-miR-17-5p were 0.724 (sensitivity 51.8%, specificity 91.4%) and 0.77 (sensitivity 84.3%, specificity 57.1%) respectively. The AUC of EMVs combined with EMV-miR-17-5p was 0.815 (sensitivity 74.7%, specificity 74.3%). Taken together, our results suggested that EMVs and EMV-miR-17-5p might be used as surrogate markers for vascular aging.

## Discussion

This study showed that y.EMVs prevented cerebrovascular aging as indicated by inhibiting vascular senescence, CBF reduction and BBB dysfunction, and alleviated brain aging including decreasing hippocampus senescent cell number, aging hallmarks (TERT, klotho, MDA, GSSG) and enhancing mice cognitive function, whereas o.EMVs played the contrary roles. The effects of EMVs were more obvious than EEXs. Mechanism study on EMVs showed that y.EMVs inhibited while o.EMVs promoted cerebrovascular aging by regulating content miR-17-5p level. Furthermore, we found that y.EMVs attenuated ECs senescence and improved ECs functions including viability, oxidative stress and apoptosis by activating the miR-17-5p/PI3K/Akt pathway. O.EMVs functioned in the opposite way. Clinically, we found that plasma EMVs level was significantly increased in healthy individuals aged over 65 and positively correlated with the vascular aging indicator IMT, while the plasma level of EMV-miR-17-5p was decreased in the elderly and negatively correlated with IMT. Taken together, our study demonstrates that young ECs could release EMVs to prevent vascular aging by ameliorating ECs senescence and dysfunction via activating miR-17-5p/PI3K/Akt pathway, and subsequently inhibit brain aging. In senescent ECs, the released EMVs switched to the opposite effects, accompanied with a significant increase of EMVs level and downregulation of miR-17-5p encapsulated in it. EMVs and their contained miR-17-5p are promising vascular aging biomarkers. Our study provides potential targets for preventing and indicating cerebrovascular and brain aging.

Vascular aging is responsible for multiple vascular diseases which have brought huge burden to health care resources [3, 4]. However, there is no clinical biomarker for vascular aging. EMVs and EEXs are increasingly accepted as important mediators and cellular biomarkers of vascular injury in many diseases, including stroke, diabetes and renal diseases [29, 30]. However, little is known about the correlation between EMVs/EEXs and vascular aging. Here, we first found that y.EMVs/y.EEXs markedly reduced while o.EMVs/o.EEXs increased the senescent cell number of basilar artery, and the role of EMVs was more significant, indicating the potential roles of EMVs/EEXs in regulating cerebrovascular aging. As we known, CBF and BBB are impaired due to the progress of cerebrovascular aging [4, 31]. One study demonstrated that the cerebral microvessel density and CBF significantly dropped by 15% and 33% respectively in the old mice [32]. Advanced dynamic contrast-enhanced magnetic resonance imaging analysis has shown that BBB permeability is obviously higher in hippocampus of older people [33]. In this study, we showed that y.EMVs and y.EEXs improved the cerebrovascular density, CBF and BBB integrity of o.mice, while o.EMVs and o.EEXs played the opposite roles in young mice. These data implied that young ECs could secrete rejuvenated EMVs/EEXs which ameliorate cerebrovascular aging, but EMVs/EEXs generated from senescent ECs accelerate cerebrovascular aging when getting old. Of interesting, the effects of EMVs were more significant than EEXs in our study. The biogenesis of EMVs and EEXs is distinct [7, 9]. Their different roles in regulating cell senescence deserve further exploration. Our findings are supported by previous observations showing that EMVs from senescent pig heart ECs induced ECs senescence [34], and EEXs from senescent HUVECs induced vascular smooth muscle cells senescence [35]. By using primarily cerebral ECs, our results first suggested that the effects of EMVs were more effective than EEXs on regulating cerebrovascular aging.

Cerebrovascular aging plays a prominent role in the pathophysiological process of brain aging [4, 31]. We further detected the effects of EMVs/EEXs on brain aging. SA-β-gal staining showed that y.EMVs/y.EEXs decreased while o.EMVs/o.EEXs increased cell senescence in mice hippocampus. Aged brain undergoes progressive mythological and functional changes resulting in the observed behavioral retrogressions, like cognitive impairment [31]. We found that y.EMVs/y.EEXs improved, while o.EMVs/o.EEXs insulted cognitive ability of mice according to the morris water test. EMVs were more effective than EEXs in the escape latency performance and searching strategy, suggesting that EMVs is superior than EEXs at alleviating brain aging. The aging markers in hippocampus revealed that y.EMVs and y.EEXs significantly reduced senescent cell number, and decreased the levels of MDA and GSSG, while increased TERT and klotho levels of old mice. O.EMVs and o.EEXs played the contrary roles in regulating the above aging markers. EMVs were more effective than EEXs on regulating GSSG and klotho. MDA and GSSG have been implicated in aging and served as oxidative stress markers, and high levels of MDA and GSSG were found in the hippocampus of aged rat [36, 37]. Klotho gene is widely accepted as an anti-aging gene with anti-oxidative stress and anti-inflammation functions, and klotho overexpression could extend life span of mice [38]. TERT could maintain telomeres length to prevent oxidative stress damage and aging [39, 40]. Therefore, our data indicated that y.EMVs alleviated whereas o.EMVs exacerbated brain aging more effectively than EEXs, which might be partly related with oxidative stress regulation. One latest study demonstrated that parabiosis of young and old mice could rejuvenate neurogenesis and cognitive function of old mice [41], suggesting substances in the circulation could be the mediators. Our data imply that the circulating y.EMVs might play a primary role in rejuvenation of old mice.

Emerging evidence has identified miRNAs as important mediators of EMVs induced biological effects [7, 9, 11]. Our miRNA sequencing results showed that miR-17-5p level was significantly decreased in o.EMVs compared to that in y.EMVs. MiR-17-5p is a pivotal miRNA involved in aging [25]. MiR-17-5p was highly expressed in embryonic stem cell-derived small extracellular vesicles which could reverse senescence-related neurogenesis dysfunction [42]. Transgenic expression of miR-17 increased mouse lifespan and decreased senescent cell number in the skin, lung, kidney and heart [25]. MiR-17-5p is downregulated in stress-induced senescence of human diploid fibroblasts and human trabecular meshwork cells [43]. MiR-17 overexpression suppresses apoptosis and promotes tube formation in HUVECs [44]. These results indicate that miR-17-5p might be important for preventing aging and ECs dysfunctions. In the present study, we found that miR-17-5p levels in young brain ECs, brain tissue and y.EMVs were significantly higher in comparison to the old ones. Y.EMVs treatment could elevate miR-17-5p levels of mice brain and senescent ECs. These results demonstrate that miR-17-5p is enriched in young brain ECs and could be encapsulated in EMVs and transferred to the targeted ECs, and that miR-17-5p is down-regulated in senescent ECs of old mice. Of note, O.EMVs might transfer some negative regulators of miR-17-5p, in addition to simply decreasing the amount of miR-17-5p, and the mechanisms of the pro-aging effect of o.EMVs needs to be further elucidated. Our functional study showed that overexpressing miR-17-5p in o.EMVs or knocking down miR-17-5p in y.EMVs significantly inhibit their deleterious or beneficial effects on cerebrovascular and brain aging, indicating that miR-17-5p is pivotal in mediating these functions of EMVs.

ECs senescence and dysfunction are early features of cerebrovascular aging and brain aging [1]. We further detected the effects and the underlying mechanisms of EMVs on ECs. Results showed that y.EMVs significantly decreased while o.EMVs increased ECs senescence, which were partly inhibited by miR-17-5p down-regulation or overexpression respectively. These results indicate that EMVs play variable roles in regulating ECs senescence during aging by modulating miR-17-5p. Oxidative stress and apoptosis are the most important activities involved in vascular aging and ECs senescence [45, 46]. Excessive oxidative stress could impair ECs functions [47], resulting in BBB disruption, brain edema, and CBF reduction [48, 49]. Besides, high level of ECs apoptosis is related with CBF decline, BBB breakdown, and vascular aging [4, 47]. We measured ECs apoptosis and the generations of ROS and NO which are important mediators in oxidative stress, and found that y.EMVs inhibited while o.EMVs increased oxidative stress and apoptosis of cultured ECs or ECs in brain tissue slices. This is as expectation, since increased oxidative stress has been demonstrated to deteriorate cell apoptosis [49]. Moreover, ECs migration, proliferation and tube formation are indispensable for angiogenesis and vascular sprouting [50]. The reduced angiogenesis and vessel density are positively related with CBF decrease [4]. In present study, we found that y.EMVs promoted the angiogenic capability of ECs, and o.EMVs functioned in the opposite way. MiR-17-5p up-regulation in o.EMVs or down-regulation in y.EMVs evidently impaired the effects of y.EMVs or o.EMVs on the above ECs functions. All these data indicate that EMVs could modulate ECs senescence and functions via regulating miR-17-5p, thus to regulate vascular aging and brain aging. In molecular pathway study, IPA analysis showed that PI3K family and Akt were closely related to miR-17-5p function. PI3K family is a multi-functional family and PI3K/Akt is a classic regulatory pathway. Our previous studies have demonstrated that PI3K/Akt pathway is involved in ROS/NO production, apoptosis, cell migration and proliferation [47, 51]. Our results showed that y.EMVs promoted while o.EMVs inhibited p-PI3K/PI3K and p-Akt/Akt levels in senescent and young ECs, respectively, and which were modulated by miR-17-5p upregulation or downregulation. PI3K inhibition or activation could significantly suppress the effects of o.EMVs or y.EMVs and miR-17-5p overexpression or knockdown on p-Akt/Akt level, as well as on ECs senescence and functions. The PI3K inhibitor LY-294002 and activator 740Y-P could inhibit but not completely reverse the effects of y.EMVs/o.EMVs and o.EMV^miR-17^/y.EMV^miR-17KO^ on ECs functions, implying that PI3K/Akt is an important signaling pathway but not the only pathway contributing to EMVs and miR-17-5p functions. Based on our observations, it is conceivable that y.EMVs and o.EMVs exerted their effects on resisting or inducing cerebrovascular and brain aging via activating or inhibiting miR-17-5p/PI3K/Akt signaling pathway.

In this study, we also found that the levels of EMVs were increased in the medium of high-passage ECs and plasma of old mice with vascular aging. Levels of miR-17-5p in o.EMVs and senescent ECs were significantly decreased compared with y.EMVs and young ECs. These data suggested that EMVs and EMVs-miR-17-5p levels might be potential biomarkers for vascular aging. Interestingly, our clinical result showed that the level of EMVs was significantly increased and the level of EMV-miR-17-5p was decreased in the aged people. These results indicated that levels of EMVs and EMV-miR-17-5p were positively or negatively correlated with chronological aging, respectively. Chronological aging does not always parallel with vascular aging. It has been clinically demonstrated that IMT could reflect vascular aging and its value higher than 0.58 indicates vascular aging in healthy subjects [1, 26-28]. Nevertheless, the limitation of IMT should be noticed that the result of IMT may vary due to skills of different investigators [52]. The indicator for vascular aging should be further explored. Blood-based biomarker is characterized with objectivity and reproducibility [53]. Besides, venous blood collection is commonplace in clinical practice. In the present study, correlation analysis showed that plasma EMVs was positively while EMV-miR-17-5p was negatively correlated with IMT. ROC analysis showed the reliable diagnostic values of EMVs and EMV-miR-17-5p for vascular aging. The diagnostic efficacy of EMV-miR-17-5p is better than EMVs, and their combination is the best. These data imply that EMVs and EMV-miR-17-5p are associated with vascular aging and these findings accord with the results of in vivo and vitro studies, raising the potential that EMVs released from young ECs could rejuvenate vascular and brain aging, and EMVs released from senescent ECs functioned the other way. EMVs and EMV-miR-17-5p could serve as promising biomarkers for vascular aging. Nevertheless, our findings need more future studies by enlarging the sample size and enrolling population from other races and latitudes.

In summary, we have demonstrated that EMVs play more pivotal roles in regulating vascular aging than EEXs. Young ECs released EMVs could alleviate vascular and brain aging by ameliorating ECs senescence and protecting ECs functions via miR-17-5p/PI3K/Akt pathway. EMVs derived from senescent ECs play the opposite roles. Plasma EMVs and their carried miR-17-5p could be efficient biomarkers for vascular aging. The present study also shed new light on the mechanisms and treatment of vascular and brain aging.

## Materials and Methods

### Isolation and culture of primary mouse brain microvascular endothelial cells

Mouse brain microvascular endothelial cells (BMECs) were prepared from 2-week-old C57BL/6 mice as previously recommended [54]. Mice was soaked in 75% ethanol to disinfect 5 min after the cervical dislocation, then mice brains were removed and placed in a 10 cm petri dish containing precooled Hank’s solution. The white matter, residual blood vessels and pia mater were removed and the cerebral cortex was retained. Brain tissue was minced and homogenized, and 10 ml Modified Eagle Medium (MEM) medium was added to the tissue suspension and centrifuge the cells at 1200 rpm for 10 min at 4°C. The supernatant was discarded and myelin was removed by centrifugation at 2600 r/min for 10 min in 22% bovine serum albumin (BSA, Sigma). The microvascular segments and cells were resuspended by using the Endothelial Cell Growth Media (ECGM) and centrifuge at 1200 r/min for 5 min. Then the BMECs were inoculated in 12-well culture dishes previously coated with type I collagen. BMECs were cultured in 5% CO_2_ incubator at 37°C with medium containing 80% Dulbecco’s Modified Eagle Medium (Gibco, cat# 10566016), 15% fatal bovine serum (Gibco, cat# 16010142), 0.05% endothelial cell growth supplements (Millipore, cat# 211F-GS), 0.1% heparin (Sigma, cat# H9267) and 1% antibiotic-antimycotic solution (100X, Invitrogen, cat# 15240096). On the second day, endothelial cell growth medium (ECGM) containing 4 μg/mL puromycin (Sigma, cat# P8833) was added to continue culture, and 72 h later it was replaced with conventional ECGM medium. The culture medium was changed every other day, and when the cells reached 90-95% fusion, 0.25% trypsin was used for passage. BMECs were seeded at a density of 1.0 × 10^4^ cells/cm^2^ for subsequent experiments and passaged when BMECs reached 90% confluence. Serial passage of cells was performed by trypsinization with trypsin-EDTA (Gibco, cat# 25200072). The purity of BMECs was controlled by immunofluorescence staining of endothelium marker CD31 (1:500, CST, USA, cat# 3528S).

### Determination of cell senescence

Cell aging is induced by passage. Senescence-associated β-galactosidase (SA-β-gal) staining was applied to determine ECs senescence. BMECs were cultured in 6-well plates with 75% confluence, then the cells were washed twice by phosphate buffered solution (PBS), fixed by Fixative Solution for 15 min at room temperature (RT), and stained by X-Gal Staining Solution for 24 h at 37 °C in dark without CO_2_. SA-β-gal staining of BMECs were viewed under an inverted microscope (Nikon ECLIPSE Ti-S, Japan). Passage 2-4 (P 2-4) and passage 15-16 (P 15-16) BMECs were defined as y.cells (SA-β-gal < 5%) and o.cells (SA-β-gal > 60%), respectively [55].

### Preparation, analysis and characterization of EMVs and EEXs

The culture media of y.cells and o.cells were collected and centrifuged at 300 rcf for 15 min and 2000 rcf for 20 min and the supernatants were further centrifuged at 20,000 rcf for 70 min to pellet EMVs. These EMVs were defined as y.EMVs and o.EMVs. The supernatants of y.EMVs and o.EMVs were further ultracentrifuged at 100,000 rcf for 2 h to pellet EEXs and defined as y.EEXs and o.EEXs, respectively. EMVs and EEXs were diluted 1/200 in PBS and their levels were analyzed by NTANS300 Instrument (Malvern Instruments, United Kingdom) under light scatter mode with a 405-nm laser. Three videos of typically 30 s duration were taken, with a frame rate of 30 frames per second. Data with corresponding standard error were analyzed by NTA 3.0 software (Malvern Instruments, United Kingdom). Absolute numbers were recorded and back-calculated using the dilution factor. The morphology and size of EMVs and EEXs were assessed by TEM as we previously described [56]. EMVs and EEXs were respectively diluted with PBS to the concentration of 2 × 10 ^8^/ml, and the dilution was added to a copper grid and incubated at RT for 5 min. Filter paper was used to absorb unevaporated solution. EMVs and EEXs were negatively stained with 2% aqueous uranylacetate at RT for 5 min and dried at 25°C for 30 min. EMVs an d EEXs were examined at 80 kV in a Hitachi JEM-1400 transmission electron microscope (Hitachi, Tokyo, Japan), and all images were recorded with Gatan832 CCD digital camera. Laboratory technicians who measured the levels of EMVs/EEXs were blinded to the information of the grouping.

### Incorporation study of EMVs and EEXs in vitro and in vivo

1×10^9^ particles/ml EMVs and EEXs were pre-incubated with PKH 26 dye (2 μM, Sigma, USA, cat# PKH26GL) [57]. Briefly, 2×10^8^/ml EMVs and EEXs were put into 2 ml EP tube (marked A) and mixed with 200 μl diluent C (Sigma, USA, cat# CGLDIL), another EP tube containing 4 μl PKH 26 and 200 μl diluent C was marked B, then tubes A and B were mixed immediately at RT for 3min, the mixture was transferred into 15 ml centrifuge tube and 2 ml 1% bovine serum albumin (BSA; Sigma, USA, cat# A4628) was added to terminate dyeing. The total mixture was transferred into ultracentrifuge tube and centrifuged at 20,000 rcf for 70 min to pellet EMVs, and the supernatants was further ultracentrifuged at 100,000 rcf for 2 h to pellet EEXs. The stained EMVs and EEXs were added into BMECs to incorporate for 8 h in a 37 °C incubator, and cells were further incubated with DIO (1:300, Beyotime, China, cat# C1038) for 20 min at RT. As for the incorporation into mouse brain vascular, 100 μl saline containing 1×10^9^ particles/ml PKH 26 stained EMVs or EEXs were injected into mice by a single bolus administration through tail vein [58]. Slice preparation were performed as previously recommended [59]. At 12 h, mice were anaesthetized by isoflurane (2–2.5% in a 70% N_2_O/30% O_2_ mixture) and transcardially perfused with 4% buffered paraformaldehyde solution (pH 7.4). The hippocampus was separated and maintained in 4% paraformaldehyde for 24 h at 4 °C, then transferred into 15% sucrose solution for 24 h at 4 °C followed by immersion in 30% sucrose for additional 24 h at 4 °C. The brains were then embedded with optimal cutting temperature compound (O.C.T, Solarbio, China) and finally stored in -80 °C. Brains were cut into coronal 15 μm thick sections with a cryostat. Sections were fixed by Immunol-staining fix solution (Beyotime, China, cat# P0098) for 30 min, then were washed three times with PBS and permeabilized with 0.1% Triton X-100 (Beyotime, China, cat# P0096) for 30 min on ice. Samples were blocked with QuickBlock™ blocking buffer (Beyotime, China, cat# P0260) for 60 min at RT, then were incubated with primary antibodies mouse monoclonal anti-CD31 (1:500, CST, USA, cat# 3528S) at 4 °C overnight, and then incubated with goat anti-mouse IgG H&L (Alexa Fluor® 647) (1:500, Abcam, cat# ab150115) for 60 min. Finally, DAPI (1 μM, Beyotime, cat# C1006) was used for staining cellular nuclear in BMECs and brain slices. Fluorescence was detected under a confocal microscope (OLYMPUS, FV3000, Japan).

### Animals, sample size calculation and grouping

All animal experiments were performed following the ARRIVE guidelines and complied with the National Institutes of Health Guide for the Care and Use of Laboratory Animals and were approved by the Institutional Animal Care and Use Committee of Guangdong Medical University (Permitted Number: GDY1902103). The C57BL/6 male and female mice were purchased from Nanjing Biomedicine Institute of Nanjing University and housed in a temperature and light-controlled room with free access to water and food. Mice with 3-month and 24-month ages were defined as young mice (y.mice) and old mice (o.mice), respectively.

Sample size calculation was determined by the Power Analysis and Sample Size 15 (PASS 15) software using “Confidence Intervals for One Variance using Variance” for predicting detectable differences to reach power of 0.80 at a significance level of <0.05 and assuming a 45% difference at the 90% confidence level. Results showed that at least 10 mice per group were needed to suffice statistically significant differences. We performed Morris water maze and measured CBF with laser doppler flowmetry in 10 mice, and the 6 of them were used for TERT, klotho and MDA analysis (brain tissue). Immunofluorescence study and β-galactosidase staining of hippocampus and β-galactosidase staining of basilar artery were also detected in these 6 mice. In another set of experiments, the plasma of 6 mice were collected and used for GSSG analysis. The rest 4 mice were accepted Evans blue (EB) extravasation analysis. Mice were randomly grouped based on their ear tag identification numbers.

Firstly, we compared the effects of y.EMVs/o.EMVs and y.EEXs/o.EEXs on mouse cerebrovascular and brain aging. Mice were randomly divided into six experimental groups (n=10/group): y.mice+saline, y.mice+o.EMVs, y.mice+o.EEXs, o.mice+saline, o.mice+y.EMVs, o.mice+y.EEXs. Mice were injected with various EMVs/EEXs with the concentration of 1×10^10^ particles/100 μl in 0.9% saline or with 100 μl 0.9% saline as control through tail vein once a week for five consecutive weeks.

### Cerebral blood flow and blood brain barrier function measurements

The CBF of mice was determined by laser doppler flowmetry under the PeriCam PSI System (Perimed, Sweden) as we previously indicated [47]. In brief, mice were anesthetized by isoflurane (2–2.5% in a 70% N_2_O/30% O_2_ mixture) and placed on a stereotaxic apparatus. A cross-cut incision was made on the skin of head to expose the entire skull. The intact skull was scanned under the PeriCam PSI System for 60 s. The CBF perfusion was analyzed by the Pimsoft software. BBB injury was detected by Evans blue (EB) extravasation after various EMVs/EEXs treatments as previously reported [60]. Briefly, 2% solution of EB (Sigma-Aldrich, cat# E2129) in saline (4 ml/kg of body weight) was injected through tail vein. Six hours later, mice were anesthetized and the organs were perfused with phosphate buffer saline (PBS) and mouse brain was isolated and weighed. Brain tissue was immersed in 50% trichloroacetic acid (Sigma-Aldrich, cat# T0699) for 48 h. Hemisphere samples were homogenized in 1ml 50% trichloroacetic acid, the homogenate was ultracentrifuged at 5000 rcf for 20 min 4 °C. Supernatant was collected, and each 60 μl supernatant was added 80 μl anhydrous ethanol. EB level in the supernatant was measured at 620 nm by using a microplate spectrophotometer (Molecular Devices, Sunnyvale, CA, USA). Additionally, the absorbance value was adjusted to the volume of EB using the standard curve. The standard curve of EB could be found in the supplementary materials. The final EB dye value was represented as ug/g tissue.

### SA-β-gal staining in mice hippocampus and basilar artery

As the basilar artery was the easiest observed vascular in mice brain, we chose to stain basilar artery with SA-β-gal kit to assess cerebrovascular aging. As previously reported [47, 61], mouse brain was placed in a 10 cm petri dish containing PBS, then the brainstem was gently separated and was placed vertically in O.C.T (Solarbio, China). Brain tissues were cut into coronal 20-μm-thick sections by a cryotome. The SA-β-gal staining was carried out according to the manufacturer’s instructions [62]. In brief, brain slices of hippocampus and basilar artery were fixed by fixative solution for 15 min at RT, and stained by X-Gal staining solution for 24 h at 37 °C in dark without CO_2_. Brain slices of hippocampus was viewed under an inverted microscope (Nikon ECLIPSE Ti-S, Japan). Brain slices of basilar artery was assessed under the Laser microdissection pressure capture system (Palm microbeam, Zeiss, Germany). The intensity of blue-stained cells and total cells in basilar artery and CA3 area of the hippocampus were counted by Image J software (NIH). Data were collected from 5 random fields per mouse, 6 individual mice in each group.

### Detection of aging associated biomarkers

Aging associated biomarkers including MDA, GSSG and klotho levels were detected according to the manufacturer’s instructions (MDA: Beyotime, cat# S0131M, GSSG: Beyotime, cat# S0053, Klotho: ZCI Bio, cat# ZC-31968) as previously reported, and TERT level was measured by q-PCR [36, 37, 39, 40]. The detailed procedures could be found in the supplementary materials.

### Morris water maze test

Cognitive ability of the mice was analyzed by Morris Water Maze task as previously described [63]. The maze was a cylindrical test apparatus (1 m in diameter) filled with water at 25±1°C. The tank was divided into 4 quadrants, and contained a circular escape platform (10 cm in diameter), to which the mice could escape. The escape platform was placed 1 cm below the surface of the water and hidden from view by making the water opaque with a white bio-safe material. Data were obtained from a video camera which was connected to an automated tracking system (Zhongshi Di Chuang Technology Development Co., Ltd., China), fixed 1.7 m above the center of the pool. There were visual cues around the water maze and remained in the same position throughout the training and testing periods. Oriented navigation trials were performed 4 times per day for 5 consecutive days with a 1 h interval between trials. Each mouse had a time limit of 60 s to find the hidden platform and then required to remain seated on the platform for 5 s. If the mouse could not find the platform, the mouse was gently guided to the platform and allowed to re-orient to the distal visual cues for an additional 10 s before being removed from the pool. At day 6, the platform was removed, and the probe trial was conducted to assess the extent of memory. Each mouse was released from the start position opposite to the former platform quadrant. They were allowed 60 s to search for the platform. Time spent in the target quadrant, the number of target crossings over the previous location of the target platform and probe strategy were recorded.

### Immunofluorescent staining of mouse brain slices

As we previously reported [47, 57], mouse brains were cut into coronal 20-μm-thick sections with a cryostat. Levels of cerebral microvascular density, and ROS/NO and apoptosis colocalized with ECs in mouse hippocampus were assessed. The sections were incubated with mouse monoclonal anti-CD31 (1:500, CST, USA, cat# 3528S) at 4 °C overnight, and then incubated with goat anti-mouse IgG H&L (Alexa Fluor® 647) (1:500, Abcam, cat# ab150115) for 60 min. Brain sections were washed three times, 8 min each time with Immunol-staining wash buffer (Beyotime, cat# P0106). After washing, the coronal slices were continued to be stained by DCFH-DA (10 μM, Beyotime, cat# S0033S) or DAF-FM DA (5 μM, Beyotime, cat# S0019) and incubated for 30 min at 37 °C out of light for ROS and NO staining. For apoptosis analysis, slices were stained with TUNEL solution (Beyotime, cat# C1086) and incubated for 60 min at 37 °C in the dark. Nuclei were counterstained with DAPI (1 μM, Beyotime, cat# C1006) for 10 min at RT. DCFH-DA and TUNEL staining were detected with an excitation wavelength of 488 nm, DAF-FM DA staining was detected with an excitation wavelength of 495 nm. The positive staining was observed under the same fluorescent microscope settings, and the percentage of positive cells of each slice was calculated from the average value of 5 randomly selected fields under a fluorescence microscope (OLYMPUS, FV3000, Japan). The average of three sequential slices from rostral to caudal represented the data for individual mouse. Images were processed with FV31S-SW (Ver 2.3.1) and analyzed by Image J (NIH) software. All slices were analyzed by a laboratory technician who was unaware of the group studied.

### MicroRNA sequencing

Total RNA of y.EMVs and o.EMVs were isolated using the miRNeasy kit (Qiagen Inc., Valencia, CA, USA) according to the manufacturer’s instructions, respectively. The concentration and integrity of the extracted total RNA was estimated by Qubit 3.0 Fluorometer (Invitrogen, Carlsbad, California) and Agilent 2100 Bioanalyzer (Applied Biosystems, Carlsbad, CA), respectively. Library preparation for small RNA sequencing was prepared with approximately 100 ng of total RNA using VAHTSTM Small RNA Library Prep Kit for Illumina® (Vazyme Biotech) [64]. For miRNA expression analysis, exceRpt was used to estimate the miRNA expression in miRBase. Alternatively, novel miRNAs were identified with miRDeep2. TMM (trimmed mean of M-values) was used to normalize the gene expression. Differentially expressed genes were identified using the edgeR program. Genes showing altered expression with *P* < 0.05 and more than 1.5-fold changes were considered differentially expressed.

### RNA extraction and quantitative reverse transcription PCR (RT-PCR)

The level of differentially expressed miR-17-5p was analyzed by RT-PCR in y.EMVs/o.EMVs, BMECs and in the mouse brain [56]. The total RNAs were extracted using TRIzol reagent (Invitrogen, Carlsbad, CA), and miRNAs in y.EMVs/o.EMVs were further extracted by miRNeasy Mini Kit under manufacturer’s instruction and the detail information could be found in the supplementary materials. The quality and concentration of RNA was detected by spectrophotometer. MiR-17-5p expression was quantified using hairpin-it TM miRNAs RT-PCR Quantitation kit (GenePharma, Shanghai, China) based on manufactory instruments (25 °C for 30 min, 42 °C for 30 min, and 85 °C for 5 min). Real-time PCR was carried out on a LightCycer480-II System (Roche Diagnostics, Penzberg Germany) by using SYBR Premix Ex Taq TM (TAKARA, Japan) and parameters were: 95 °C for 3 min, 40 cycles were performed at 95 °C for 12 s, 60 °C for 40 s. Mmu-miR-17-5p forward: 5’-CAA AGU GCU UAC AGU GCA GGU AG-3’, reverse: 5’-ACC UGC ACU GUA AGC ACU UU GUU-3’. U6 was chosen as housekeeping gene for normalizing the data of miR-17-5p expression. Each experiment was repeated three times. The relative quantification of the gene expression was determined using the comparative CT method (2^-ΔΔCt^).

### MiR-17-5p overexpression and knockdown in BMECs and the grouping of experiments

Stable overexpression and knockdown of miR-17-5p in BMECs were established by using lentivirus infection method [47]. Lentiviruses expressing green fluorescent protein marker with miR-17-5p mimic (Lv-miR-17-5p), miR-17-5p silencing short hairpin RNA (shRNA) (Lv-si-miR-17-5p) or their scrambled controls (Lv-SC) were purchased from GenePharma Biotech (Shanghai, China). Lv-miR-17-5p and Lv-miR-17-5pSC were transfected into in o.cells, and Lv-si-miR-17-5p and Lv-si-miR-17-5pSC were transfected into y.cells, respectively (referred to o.cells^miR-17^, o.cells^miR-17SC^, y.cells^miR-17KO^, y.cells^miR-17KOSC^) [47]. Various EMVs were collected from the culture media of these cells, and were defined as o.EMV^miR-17^, o.EMV^miR-17SC^, y.EMV^miR-17KO^, and y.EMV^miR-17KOSC^. The levels of miR-17-5p in various cells and EMVs were confirmed by RT-PCR as we mentioned above [47]. The various EMVs were applied for animal and BMECs functional studies, and the experimental groups were as follows: y.mice/y.cells, y.mice/y.cells+o.EMVs, y.mice/y.cells+o.EMV^miR-17SC^, y.mice/y.cells+o.EMV^miR-17^, o.mice/o.cells, o.mice/o.cells+y.EMVs, o.mice/o.cells+y.EMV^miR-17KOSC^, o.mice/o.cells+y.EMV^miR-17KO^. The animal study including cerebrovascular and brain aging experiments were launched as we illustrated above.

### Analysis of BMECs senescence and functions

BMECs were cultured in 6-well plates or 96-well plates with 75% confluence and incubated with different EMVs (1×10^9^ particles/ml) for 24 h before various experiments were launched [58]. A SA-β-gal staining kit was used to assess BMECs senescence as we mentioned above. Levels of ROS/NO and apoptosis of BMECs were measured as previously reported [47, 65]. For ROS and NO analysis, BMECs were grown to confluence on 6-well cell culture plate, then incubated with Dihydroethidium (DHE, 5μM, Beyotime, China, cat# S0063) or DAF-FM DA (5μM, Beyotime, China, cat# S0019) solution at 37 °C for 1 h and 30 min, respectively, and washed three times with PBS. For apoptosis measurement, cells were fixed, washed with PBS and stained with Hoechast 33258 staining solution (Beyotime, China, cat# C0003) for 10 min. The positive cells were observed under an inverted microscope (EVOS FL AUTO, Life Technologies, USA). Five independent fields were assessed for each well, and the average number of positive cells and negative total cells per field were determined. The ROS, NO and apoptosis production of cells was defined as the ratios of positive cells in versus total cells.

Migration, proliferative capabilities and tube formation of BMECs were tested as previously reported [47, 66]. Proliferative capability of BMECs was tested by 3-[4,5-dimethylthiazyol-2-yl] -2,5-diphenyltetrazolium bromide (MTT, 5mg/ml, Sigma, cat# CT01) assay. BMECs were respectively seeded at 2 ×10^3^/96-well plate and cultured in 100 μl DMEM (Gibco, USA) supplemented with 10% Fetal bovine serum (FBS, Gibco, USA). After 2 days of incubation, the cells were treated with 20 μl MTT solution for 4 h at 37 °C, then150 μl DMSO was added to each well and cells were further incubated for 20 min at 37 °C. The optical density of the cells was read at 490 nm in a microplate reader (BioTek) in triplicate. Results were calculated from the values obtained in three independent experiments. The migration capacities were measured by scratch assay. BMECs were grown to confluence on 6-well cell culture plate. A scratch was made through the cell monolayer using a 200 μl pipette tip. After washing with PBS, cells were cultured in 0.5% FBS maintenance medium. Photographs of the wounded area were taken immediately (0-h time point) and 16 h after making the scratch to monitor the invasion of cells into the scratched area. The tube formation ability was evaluated by using the tube formation assay kit (Chemicon, USA, cat# ECM625) according to the manufacturer’s instructions. Briefly, ECMatrix working solution was thawed on ice overnight, mixed with 10×ECMatrix diluents and placed in a 96-well tissue culture plate at 37°C for 1h to allow the matrix solution to solidify. BMECs were replaced (1×10^4^ cells/well) on the surface of the solidified ECMatrix and incubated with endothelial cell growth medium-2 (EGM-2) (Longza, USA) for 24 h at 37°C. Various EMVs (1×10^9^ particles/ml) were added in each well and the cells were further incubated for 24 h at 37 °C, then tube formation was evaluated with an inverted microscope (EVOS FL AUTO, Life Technologies, USA). A structure exhibiting a length 3 times its width was defined as a tube. Five independent fields were assessed for each well, and the average number of tubes per field was determined.

### MiR-17-5p regulatory network analyzed by Ingenuity Pathway Analysis

The miR-17-5p regulatory network was analyzed by using IPA (Ingenuity Systems; Mountain View, CA, USA), which assists with RNA sequencing via grouping of differentially expressed genes into known functions, pathways, and networks primarily based on human and rodent studies [67]. The identified genes were mapped to genetic networks available from the Ingenuity database and were then ranked by score. The significance was set at a *P* < 0.05. Based on the results of IPA analysis, for molecular mechanism study, we pretreated y.cells with PI3K activator (740Y-P, 10 μM; Selleckchem), and pretreated o.cells with PI3K inhibitor (LY294002, 20 μM; Selleckchem) for 2 h before functional in vitro studies were launched.

### Western blot analysis

Total 25 μg proteins of ECs were extracted with cell lysis buffer (Applygen Technologies Company, China, cat# C1051-100) supplemented with protease inhibitor tablet (Thermo Scientific, USA, cat# 36978). Proteins were loaded into a 12% SDS-PAGE gels and transferred onto polyvinylidene difluoride membranes (pore size: 0.22 mm, Millipore, Billerica, USA). The membranes were blocked with 5% non-fat milk for 1 h and incubated with primary antibodies against β-actin (1:1000, Abcam, cat# ab8226), PI3K (1:1000, CST, USA, cat# 4249S), phospho-PI3K (1:1000, CST, USA, cat# 17366S), Akt (1:1000, CST, USA, cat# 4685S) and phospho-Akt (1:1000, CST, USA, cat# 4060S) overnight at 4 °C. The secondary antibody goat anti-rabbit IgG H&L (1:5000, Abcam, cat# ab6721) was added onto the membrane [47]. Chemiluminescence was assessed by Azure C600 (Azure Biosystems, America).

### Enrollment of participants

The human study was in accordance with the ethical standards of the institutional and/or national research committee and with the 1964 Helsinki declaration and its later amendments or comparable ethical standards, which were approved and conducted under supervision of ethics committee of Affiliated Hospital of Guangdong Medical University (Human Investigation Committee PJ2019-046). The trial registration number is ChiCTR2000032734 (https://www.chictr.org.cn/).

Participants were recruited from the Affiliated Hospital of Guangdong Medical University from January 2019 to December 2020 and total 119 Chinese participants were enrolled. Power Analysis and Sample Size (PASS) software was used to calculate the sample size (provided in supplementary materials). Written informed consent was obtained from all participants. Inclusion criteria were: (1) male or female aged >18 years old. (2) health in physical examination. Exclusion criteria: (1) subjects with coronary heart disease, congestive heart failure, hypertension (arterial blood pressure ≥140/90 mmHg), peripheral artery disease, stroke, tumor, pulmonary diseases, diabetes, fasting glucose ≥ 7.0 mM, osteoporosis, renal failures, cancer, dementia, and depression. (2) pregnant woman or breast-feeding woman. 119 subjects were recruited and assigned into three groups, young group (18-45 years old, n = 58), middle-age group (46–65 years old, n = 33), aged group (> 65 years old, n = 28) [68, 69].

### Intima-media thickness measurement

The Intima-media thickness (IMT) was examined by carotid ultrasound with a head elevation of 45° and a side tilt of 30°to the right in the supine position, and then to the left, the mean value was calculated [70]. Measurements were performed three times by a single investigator who was blinded to the clinical characteristics and categorization of the participants. IMT > 0.58 mm indicates vascular aging [1, 26-28].

### Analysis of the levels of EMVs and miR-17-5p in EMVs from plasma

Plasma EMVs was isolated using available kits (Life technology and MiltenyiBiotec) as we previously described [56]. In brief, 8 ml of vein blood samples were collected in ethylene diamine tetraacetic acid (EDTA)-anticoagulant tubes and were centrifuged at 1500 rcf for 5 min at 10 °C to remove blood cells and collect plasma. Then the supernatant was centrifuged at 300 rcf for 15 min and 2000 rcf for 20 min and further centrifuged at 20,000 rcf for 70 min to pellet MVs. We defined EMVs as CD105^+^CD144^+^. Samples were incubated with Annexin V antibody (1:200 dilution; Santa Cruz Biotechnology Cat# sc-1929, RRID: AB_2274151) for 2 h followed by incubation with rabbit anti-goat IgG conjugated with Q-dot®655 (1:350 dilution; Life Technologies) for 90 min at room temperature, then added with PBS supplemented with 2 mM calcium solution to give a final volume of 700 μl. The pelleted MVs were incubated with 10 μl of Biotin-conjugated anti-CD105 (Miltenyi Biotec cat# 130-094-916, RRID: AB_10828348) followed by adding 10 μl of anti-Biotin microbeads (MiltenyiBiotec cat# 130-094-256, RRID: AB_244366). Then the microbeads-labeled MVs from the total MVs suspension were separated by using a DynaMag-2 magnet (Life technology). After an overnight magnetic separation, the microbeads-bound MVs were resuspended with 100 μl particle-free PBS. 10 μl of multisort release reagent (MiltenyiBiotec) was added to each sample to cleave off the microbeads. After an overnight reaction, the MVs in the fluid were collected and considered as CD105^+^MVs. The isolated CD105^+^MVs were incubated with antibody against CD144^+^ at 4 °C overnight (1:200 dilution; Santa Cruz Biotechnology), followed by incubation with rabbit anti-goat IgG conjugated with Q-dot 655 for 2 h at 4 °C (1:350 dilution; Life Technologies). Then, 700 μl filtered PBS was added to the MV suspension. The Q-dot 655-labeled MVs were considered as CD105^+^CD144^+^ MVs. Each sample was analyzed in triplicate.

For detecting the EMV-miR-17-5p in human plasma, TRIzol reagent (Invitrogen, Carlsbad, CA) and miRNeasy Mini Kit were applied to extract miRNAs from EMVs under manufacturer’s instruction. We used quantitative reverse transcription-PCR (qRT-PCR) to determine the level of EMV-miR-17-5p. The cDNAs were synthesized from 20 μg total RNA by hairpinit TM miRNAs RT-PCR Quantitation kit (GenePharma, Shanghai, China). Hsa miR-17-5p Forward primer: 5’-GGG GCA AAG TGC TTA CAG TG-3’. Reverse primer: 5’-GTG CGT GTC GTG GAG TCG-3’. U6 Forward primer: 5’-GCT TCG GCA GCA CAT ATA CTA AAA T-3’, Reverse primer: 5’-CGC TTC ACG AAT TTG CGT GTC AT-3’. U6 was chosen as housekeeping gene for normalizing the data of miR-17-5p expression. Each experiment was repeated three times. Relative expression level of EMV-miR-17-5p was calculated using 2^-ΔΔCt^ method.

### Statistical analysis

SPSS software version 22.0 and GraphPad Prism 5 software were used for statistical analysis. D’Agostino-Pearson omnibus test was used to determine normal distribution. Comparisons for two groups with normal distribution were examined by Student’s *t*-test. Multiple comparisons were performed by one or two-way ANOVA. Correlations between variables were analyzed by Spearman correlation analysis. Results were represented as adjusted odds ratios (OR) with the corresponding 95% confidence intervals (95% CI). ROC analysis and the AUC were applied to determine the sensitivity and specificity of the EMVs, EMV-miR-17-5p and their combination for assessment of vascular aging. All data was presented as means ± SEM. For all tests, *ns*, no significant; *, *P* < 0.05; **, *P* < 0.01; ***, *P* < 0.001. For mouse and clinical experiments, ‘*n*’ represents the number of mice used and the enrolled number.

## Data Availability Statement

All relevant data are within the paper and its Supporting Information files.

## Acknowledgements

We thank Professor Yanfang Chen at Wright State University for read proof of our manuscript.

## Abbreviations

ECs: endothelial cells
EMVs: endothelial microvesicles
EEXs: endothelial exosomes
HUVECs: human umbilical vein endothelial cells
BMECs: brain microvascular endothelial cells
SA-β-gal: senescence-associated β-galactosidase
CBF: cerebral blood flow
BBB: blood brain barrier
GSSG: oxidized glutathione
MDA: malondialdehyde
TERT: telomerase reverse transcriptase
cMVD: cerebral microvascular density
IMT: intima-media thickness
DHE: Dihydroethidium
MTT: 3-[4,5-dimethylthiazyol-2-yl] -2,5-diphenyltetrazolium bromide
qRT-PCR: quantitative reverse transcription PCR
FBS: Fetal bovine serum
EGM-2: endothelial cell growth medium-2
IPA: Ingenuity Pathway Analysis

## Author Contributions

**Conceptualization:** Huiting Zhang, Zi Xie, Xiaotang Ma, Bin Zhao.

**Data curation:** Huiting Zhang, Zi Xie, Bin Zhao.

**Formal analysis:** Huiting Zhang, Zi Xie.

**Funding acquisition:** Huiting Zhang, Xiaotang Ma.

**Investigation:** Huiting Zhang, Zi Xie, Yuhui Zhao, Yanyu Chen, Xiaobing Xu, Ye Ye, Yuping Yang, Wangtao Zhong.

**Methodology:** Huiting Zhang, Zi Xie.

**Software:** Huiting Zhang, Zi Xie.

**Supervision:** Huiting Zhang, Xiaotang Ma, Bin Zhao.

**Validation:** Huiting Zhang.

**Visualization:** Huiting Zhang, Zi Xie, Yanyu Chen.

**Writing – original draft:** Huiting Zhang.

**Writing – review & editing:** Huiting Zhang, Xiaotang Ma, Bin Zhao.

## Supporting information

**S1 Fig. The analysis of EMVs and miR-17-5p levels**. (A) Plasma EMVs levels of y.mice and o.mice detected by NTA (Student’s t-test, mean ± SEM, n = 6 technical replicates, ^**^ *p* < 0.01). (B) EMVs levels of y.cells and o.cells detected by NTA (Student’s *t*-test, mean ± SEM, n = 6 technical replicates, ^***^ *p* < 0.001). (C, D) MiR-17-5p levels in various cell groups (Student’s *t*-test, mean ± SEM, n = 6 technical replicates, ^***^ *p* < 0.001). (E, F) MiR-17-5p levels in various EMVs groups (one-way ANOVA, Tukey’s multiple comparisons test, mean ± SEM, n =6 technical replicates, ^***^ *p* < 0.001).

**S2 Fig. NTA analysis of plasma EMVs from young, middle-age and aged participants**. (A-C) Representative plots showing the size distribution and concentration of plasma EMVs isolated from young, middle-age and aged participants. The CD105^+^ beads isolated MVs under fluorescence/non-fluorescence modes. Orange curve: CD105^+^MVs were measured under light scatter (non-fluorescence) mode. Red curve: CD105^+^CD144^+^Q-dots MVs were measured under fluorescence mode. n = 4 technical replicates.

**S1 Table. Molecules involved in miR-17-5p regulatory network**.

**S2 Table. Character of the enrolled patients**.

**S3 Table. AUC, sensitivity, specificity, cut-off levels, and Youden Index for EMVs and EMV-miR-17-5p in vascular aging**.

## References

1. Hamczyk MR, Nevado RM, Barettino A, Fuster V, Andrés V. Biological Versus Chronological Aging: JACC Focus Seminar. J Am Coll Cardiol. 2020;75(8):919–30. https://doi.org/10.1016/j.jacc.2019.11.062 PMID: 32130928

2. Van Skike CE, Lin AL, Roberts Burbank R, Halloran JJ, Hernandez SF, Cuvillier J, et al. mTOR drives cerebrovascular, synaptic, and cognitive dysfunction in normative aging. Aging Cell. 2020;19(1):e13057. https://doi.org/10.1111/acel.13057 PMID: 31693798

3. Katan M, Luft A. Global Burden of Stroke. Semin Neurol. 2018;38(2):208–11. https://doi.org/10.1055/s-0038-1649503 PMID: 29791947

4. Yang T, Sun Y, Lu Z, Leak RK, Zhang F. The impact of cerebrovascular aging on vascular cognitive impairment and dementia. Ageing Res Rev. 2017;34:15–29. https://doi.org/10.1016/j.arr.2016.09.007 PMID: 27693240

5. Rossman MJ, Kaplon RE, Hill SD, McNamara MN, Santos-Parker JR, Pierce GL, et al. Endothelial cell senescence with aging in healthy humans: prevention by habitual exercise and relation to vascular endothelial function. Am J Physiol Heart Circ Physiol. 2017;313(5):H890–h5. https://doi.org/10.1152/ajpheart.00416.2017 PMID: 28971843

6. Liberale L, Montecucco F, Tardif JC, Libby P, Camici GG. Inflamm-ageing: the role of inflammation in age-dependent cardiovascular disease. Eur Heart J. 2020;41(31):2974–82. https://doi.org/10.1093/eurheartj/ehz961 PMID: 32006431

7. Yin Y, Chen H, Wang Y, Zhang L, Wang X. Roles of extracellular vesicles in the aging microenvironment and age-related diseases. J Extracell Vesicles. 2021;10(12):e12154. https://doi.org/10.1002/jev2.12154 PMID: 34609061

8. Fruhbeis C, Kuo-Elsner WP, Muller C, Barth K, Peris L, Tenzer S, et al. Oligodendrocytes support axonal transport and maintenance via exosome secretion. PLoS Biol. 2020;18(12):e3000621. https://doi.org/10.1371/journal.pbio.3000621 PMID: 33351792

9. Zietzer A, Hosen MR, Wang H, Goody PR, Sylvester M, Latz E, et al. The RNA-binding protein hnRNPU regulates the sorting of microRNA-30c-5p into large extracellular vesicles. J Extracell Vesicles. 2020;9(1):1786967. https://doi.org/10.1080/20013078.2020.1786967 PMID: 32944175

10. Fruhbeis C, Frohlich D, Kuo WP, Amphornrat J, Thilemann S, Saab AS, et al. Neurotransmitter-triggered transfer of exosomes mediates oligodendrocyte-neuron communication. PLoS Biol. 2013;11(7):e1001604. https://doi.org/10.1371/journal.pbio.1001604 PMID: 23874151

11. Bei Y, Das S, Rodosthenous R, Holvoet P, Vanhaverbeke M, Monteiro M, et al. Extracellular Vesicles in Cardiovascular Theranostics. Theranostics. 2017;7(17):4168–82. https://doi.org/10.7150/thno.21274 PMID: 29158817

12. Ma X, Wang J, Li J, Ma C, Chen S, Lei W, et al. Loading MiR-210 in Endothelial Progenitor Cells Derived Exosomes Boosts Their Beneficial Effects on Hypoxia/Reoxygeneation-Injured Human Endothelial Cells via Protecting Mitochondrial Function. Cell Physiol Biochem. 2018;46(2):664–75. https://doi.org/10.1159/000488635 PMID: 29621777

13. Pan Q, Ma C, Wang Y, Wang J, Zheng J, Du D, et al. Microvesicles-mediated communication between endothelial cells modulates, endothelial survival, and angiogenic function via transferring of miR-125a-5p. J Cell Biochem. 2019;120(3):3160–72. https://doi.org/10.1002/jcb.27581 PMID: 30272818

14. Bammert TD, Hijmans JG, Reiakvam WR, Levy MV, Brewster LM, Goldthwaite ZA, et al. High glucose derived endothelial microparticles increase active caspase-3 and reduce microRNA-Let-7a expression in endothelial cells. Biochem Biophys Res Commun. 2017;493(2):1026–9. https://doi.org/10.1016/j.bbrc.2017.09.098 PMID: 28942148

15. Chung J, Kim KH, Yu N, An SH, Lee S, Kwon K. Fluid Shear Stress Regulates the Landscape of microRNAs in Endothelial Cell-Derived Small Extracellular Vesicles and Modulates the Function of Endothelial Cells. Int J Mol Sci. 2022;23(3). https://doi.org/10.3390/ijms23031314 PMID: 35163238

16. Gröne M, Sansone R, Höffken P, Horn P, Rodriguez-Mateos A, Schroeter H, et al. Cocoa Flavanols Improve Endothelial Functional Integrity in Healthy Young and Elderly Subjects. J Agric Food Chem. 2020;68(7):1871–6. https://doi.org/10.1021/acs.jafc.9b02251 PMID: 31294557

17. Alique M, Ruíz-Torres MP, Bodega G, Noci MV, Troyano N, Bohórquez L, et al. Microvesicles from the plasma of elderly subjects and from senescent endothelial cells promote vascular calcification. Aging (Albany NY). 2017;9(3):778–89. https://doi.org/10.18632/aging.101191 PMID: 28278131

18. Riquelme JA, Takov K, Santiago-Fernández C, Rossello X, Lavandero S, Yellon DM, et al. Increased production of functional small extracellular vesicles in senescent endothelial cells. J Cell Mol Med. 2020;24(8):4871–6. https://doi.org/10.1111/jcmm.15047 PMID: 32101370

19. Zengin A, Jarjou LM, Janha RE, Prentice A, Cooper C, Ebeling PR, et al. Sex-Specific Associations Between Cardiac Workload, Peripheral Vascular Calcification, and Bone Mineral Density: The Gambian Bone and Muscle Aging Study. J Bone Miner Res. 2021;36(2):227–35. https://doi.org/10.1002/jbmr.4196 PMID: 33118663

20. Villeda SA, Plambeck KE, Middeldorp J, Castellano JM, Mosher KI, Luo J, et al. Young blood reverses age-related impairments in cognitive function and synaptic plasticity in mice. Nat Med. 2014;20(6):659–63. https://doi.org/10.1038/nm.3569 PMID: 24793238

21. Yan Y, Wu R, Bo Y, Zhang M, Chen Y, Wang X, et al. Induced pluripotent stem cells-derived microvesicles accelerate deep second-degree burn wound healing in mice through miR-16-5p-mediated promotion of keratinocytes migration. Theranostics. 2020;10(22):9970–83. https://doi.org/10.7150/thno.46639 PMID: 32929328

22. Jansen F, Yang X, Hoelscher M, Cattelan A, Schmitz T, Proebsting S, et al. Endothelial microparticle-mediated transfer of MicroRNA-126 promotes vascular endothelial cell repair via SPRED1 and is abrogated in glucose-damaged endothelial microparticles. Circulation. 2013;128(18):2026–38. https://doi.org/10.1161/circulationaha.113.001720 PMID: 24014835

23. Gollmann-Tepeköylü C, Pölzl L, Graber M, Hirsch J, Nägele F, Lobenwein D, et al. miR-19a-3p containing exosomes improve function of ischaemic myocardium upon shock wave therapy. Cardiovasc Res. 2020;116(6):1226–36. https://doi.org/10.1093/cvr/cvz209 PMID: 31410448

24. Greenberg SM. Vascular Contributions to Brain Health: Cross-Cutting Themes. Stroke. 2022;53(2):391–3. https://doi.org/10.1161/strokeaha.121.034921 PMID: 35000428

25. Du WW, Yang W, Fang L, Xuan J, Li H, Khorshidi A, et al. miR-17 extends mouse lifespan by inhibiting senescence signaling mediated by MKP7. Cell Death Dis. 2014;5(7):e1355. https://doi.org/10.1038/cddis.2014.305 PMID: 25077541

26. Andreassi MG, Piccaluga E, Gargani L, Sabatino L, Borghini A, Faita F, et al. Subclinical carotid atherosclerosis and early vascular aging from long-term low-dose ionizing radiation exposure: a genetic, telomere, and vascular ultrasound study in cardiac catheterization laboratory staff. JACC Cardiovasc Interv. 2015;8(4):616–27. https://doi.org/10.1016/j.jcin.2014.12.233 PMID: 25907089

27. Engelen L, Ferreira I, Stehouwer CD, Boutouyrie P, Laurent S. Reference intervals for common carotid intima-media thickness measured with echotracking: relation with risk factors. Eur Heart J. 2013;34(30):2368–80. https://doi.org/10.1093/eurheartj/ehs380 PMID: 23186808

28. van Sloten TT, Boutouyrie P, Lisan Q, Tafflet M, Thomas F, Guibout C, et al. Body Silhouette Trajectories Across the Lifespan and Vascular Aging. Hypertension. 2018;72(5):1095–102. https://doi.org/10.1161/hypertensionaha.118.11442 PMID: 30354814

29. Otero-Ortega L, Laso-García F, Gómez-de Frutos M, Fuentes B, Diekhorst L, Díez-Tejedor E, et al. Role of Exosomes as a Treatment and Potential Biomarker for Stroke. Transl Stroke Res. 2019;10(3):241–9. https://doi.org/10.1007/s12975-018-0654-7 PMID: 30105420

30. Rodrigues KF, Pietrani NT, Fernandes AP, Bosco AA, de Sousa MCR, de Fátima Oliveira Silva I, et al. Circulating microparticles levels are increased in patients with diabetic kidney disease: A case-control research. Clin Chim Acta. 2018;479:48–55. https://doi.org/10.1016/j.cca.2017.12.048 PMID: 29305843

31. Cortes-Canteli M, Iadecola C. Alzheimer’s Disease and Vascular Aging: JACC Focus Seminar. J Am Coll Cardiol. 2020;75(8):942–51. https://doi.org/10.1016/j.jacc.2019.10.062 PMID: 32130930

32. Li Y, Choi WJ, Wei W, Song S, Zhang Q, Liu J, et al. Aging-associated changes in cerebral vasculature and blood flow as determined by quantitative optical coherence tomography angiography. Neurobiol Aging. 2018;70:148–59. https://doi.org/10.1016/j.neurobiolaging.2018.06.017 PMID: 30007164

33. Montagne A, Barnes SR, Sweeney MD, Halliday MR, Sagare AP, Zhao Z, et al. Blood-brain barrier breakdown in the aging human hippocampus. Neuron. 2015;85(2):296–302. https://doi.org/10.1016/j.neuron.2014.12.032 PMID: 25611508

34. Abbas M, Jesel L, Auger C, Amoura L, Messas N, Manin G, et al. Endothelial Microparticles From Acute Coronary Syndrome Patients Induce Premature Coronary Artery Endothelial Cell Aging and Thrombogenicity: Role of the Ang II/AT1 Receptor/NADPH Oxidase-Mediated Activation of MAPKs and PI3-Kinase Pathways. Circulation. 2017;135(3):280–96. https://doi.org/10.1161/circulationaha.116.017513 PMID: 27821539

35. Lin X, Li S, Wang YJ, Wang Y, Zhong JY, He JY, et al. Exosomal Notch3 from high glucose-stimulated endothelial cells regulates vascular smooth muscle cells calcification/aging. Life Sci. 2019;232:116582. https://doi.org/10.1016/j.lfs.2019.116582 PMID: 31220525

36. Almeida EB, Santos JMB, Paixão V, Amaral JB, Foster R, Sperandio A, et al. L-Glutamine Supplementation Improves the Benefits of Combined-Exercise Training on Oral Redox Balance and Inflammatory Status in Elderly Individuals. Oxid Med Cell Longev. 2020;2020:2852181. https://doi.org/10.1155/2020/2852181 PMID: 32411324

37. Szlęzak D, Bronowicka-Adamska P, Hutsch T, Ufnal M, Wróbel M. Hypertension and Aging Affect Liver Sulfur Metabolism in Rats. Cells. 2021;10(5). https://doi.org/10.3390/cells10051238 PMID: 34069923

38. Kurosu H, Yamamoto M, Clark JD, Pastor JV, Nandi A, Gurnani P, et al. Suppression of aging in mice by the hormone Klotho. Science. 2005;309(5742):1829–33. https://doi.org/10.1126/science.1112766 PMID: 16123266

39. Baeeri M, Didari T, Khalid M, Mohammadi-Nejad S, Daghighi SM, Farhadi R, et al. Molecular Evidence of the Inhibitory Potential of Melatonin against NaAsO(2)-Induced Aging in Male Rats. Molecules. 2021;26(21). https://doi.org/10.3390/molecules26216603 PMID: 34771016

40. Li J, Liu H, Zhao J, Chen H. Telomerase reverse transcriptase (TERT) promotes neurogenesis after hypoxic-ischemic brain damage in neonatal rats. Neurological research. 2022;44(9):819–29. https://doi.org/10.1080/01616412.2022.2056339 PMID: 35400306

41. Katsimpardi L, Litterman NK, Schein PA, Miller CM, Loffredo FS, Wojtkiewicz GR, et al. Vascular and neurogenic rejuvenation of the aging mouse brain by young systemic factors. Science. 2014;344(6184):630–4. https://doi.org/10.1126/science.1251141 PMID: 24797482

42. Hu G, Xia Y, Zhang J, Chen Y, Yuan J, Niu X, et al. ESC-sEVs Rejuvenate Senescent Hippocampal NSCs by Activating Lysosomes to Improve Cognitive Dysfunction in Vascular Dementia. Adv Sci (Weinh). 2020;7(10):1903330. https://doi.org/10.1002/advs.201903330 PMID: 32440476

43. Li G, Luna C, Qiu J, Epstein DL, Gonzalez P. Alterations in microRNA expression in stress-induced cellular senescence. Mech Ageing Dev. 2009;130(11-12):731–41. https://doi.org/10.1016/j.mad.2009.09.002 PMID: 19782699

44. Chen Z, Pan X, Sheng Z, Yan G, Chen L, Ma G. miR-17 regulates the proliferation and apoptosis of endothelial cells in coronary heart disease via targeting insulin-like-growth factor 1. Pathol Res Pract. 2019;215(9):152512. https://doi.org/10.1016/j.prp.2019.152512 PMID: 31296440

45. Huo J, Xu Z, Hosoe K, Kubo H, Miyahara H, Dai J, et al. Coenzyme Q10 Prevents Senescence and Dysfunction Caused by Oxidative Stress in Vascular Endothelial Cells. Oxid Med Cell Longev. 2018;2018:3181759. https://doi.org/10.1155/2018/3181759 PMID: 30116476

46. Miyauchi H, Minamino T, Tateno K, Kunieda T, Toko H, Komuro I. Akt negatively regulates the in vitro lifespan of human endothelial cells via a p53/p21-dependent pathway. Embo j. 2004;23(1):212–20. https://doi.org/10.1038/sj.emboj.7600045 PMID: 14713953

47. Zhang H, Pan Q, Xie Z, Chen Y, Wang J, Bihl J, et al. Implication of MicroRNA503 in Brain Endothelial Cell Function and Ischemic Stroke. Transl Stroke Res. 2020;11(5):1148–64. https://doi.org/10.1007/s12975-020-00794-0 PMID: 32285355

48. An L, Shen Y, Chopp M, Zacharek A, Venkat P, Chen Z, et al. Deficiency of Endothelial Nitric Oxide Synthase (eNOS) Exacerbates Brain Damage and Cognitive Deficit in A Mouse Model of Vascular Dementia. Aging Dis. 2021;12(3):732–46. https://doi.org/10.14336/ad.2020.0523 PMID: 34094639

49. Hu Y, Bi Y, Yao D, Wang P, Li Y. Omi/HtrA2 Protease Associated Cell Apoptosis Participates in Blood-Brain Barrier Dysfunction. Front Mol Neurosci. 2019;12:48. https://doi.org/10.3389/fnmol.2019.00048 PMID: 30853894

50. Ke Y, Li D, Zhao M, Liu C, Liu J, Zeng A, et al. Gut flora-dependent metabolite Trimethylamine-N-oxide accelerates endothelial cell senescence and vascular aging through oxidative stress. Free Radic Biol Med. 2018;116:88–100. https://doi.org/10.1016/j.freeradbiomed.2018.01.007 PMID: 29325896

51. Pan Q, Zheng J, Du D, Liao X, Ma C, Yang Y, et al. MicroRNA-126 Priming Enhances Functions of Endothelial Progenitor Cells under Physiological and Hypoxic Conditions and Their Therapeutic Efficacy in Cerebral Ischemic Damage. Stem Cells Int. 2018;2018:2912347. https://doi.org/10.1155/2018/2912347 PMID: 29760722

52. Jurasic MJ, Lovrencic-Huzjan A, Bedekovic MR, Demarin V. How to monitor vascular aging with an ultrasound. J Neurol Sci. 2007;257(1-2):139–42. https://doi.org/10.1016/j.jns.2007.01.027 PMID: 17320112

53. Montellano FA, Ungethüm K, Ramiro L, Nacu A, Hellwig S, Fluri F, et al. Role of Blood-Based Biomarkers in Ischemic Stroke Prognosis: A Systematic Review. Stroke. 2021;52(2):543–51. https://doi.org/10.1161/strokeaha.120.029232 PMID: 33430636

54. Bittner S, Ruck T, Schuhmann MK, Herrmann AM, Moha ou Maati H, Bobak N, et al. Endothelial TWIK-related potassium channel-1 (TREK1) regulates immune-cell trafficking into the CNS. Nat Med. 2013;19(9):1161–5. https://doi.org/10.1038/nm.3303 PMID: 23933981

55. Dubinsky AN, Dastidar SG, Hsu CL, Zahra R, Djakovic SN, Duarte S, et al. Let-7 coordinately suppresses components of the amino acid sensing pathway to repress mTORC1 and induce autophagy. Cell Metab. 2014;20(4):626–38. https://doi.org/10.1016/j.cmet.2014.09.001 PMID: 25295787

56. Zhang H, Chen G, Qiu W, Pan Q, Chen Y, Chen Y, et al. Plasma endothelial microvesicles and their carrying miRNA-155 serve as biomarkers for ischemic stroke. J Neurosci Res. 2020;98(11):2290–301. https://doi.org/10.1002/jnr.24696 PMID: 32725652

57. Pan Q, Kuang X, Cai S, Wang X, Du D, Wang J, et al. miR-132-3p priming enhances the effects of mesenchymal stromal cell-derived exosomes on ameliorating brain ischemic injury. Stem Cell Res Ther. 2020;11(1):260. https://doi.org/10.1186/s13287-020-01761-0 PMID: 32600449

58. Lázaro-Ibáñez E, Neuvonen M, Takatalo M, Thanigai Arasu U, Capasso C, Cerullo V, et al. Metastatic state of parent cells influences the uptake and functionality of prostate cancer cell-derived extracellular vesicles. J Extracell Vesicles. 2017;6(1):1354645. https://doi.org/10.1080/20013078.2017.1354645 PMID: 28819549

59. Chastagner P, Loria F, Vargas J, Tois J, I Diamond M, Okafo G, et al. Fate and propagation of endogenously formed Tau aggregates in neuronal cells. EMBO molecular medicine. 2020;12(12):e12025. https://doi.org/10.15252/emmm.202012025 PMID: 33179866

60. Yamazaki Y, Baker DJ, Tachibana M, Liu CC, van Deursen JM, Brott TG, et al. Vascular Cell Senescence Contributes to Blood-Brain Barrier Breakdown. Stroke. 2016;47(4):1068–77. https://doi.org/10.1161/strokeaha.115.010835 PMID: 26883501

61. Zheng LY, Li L, Ma MM, Liu Y, Wang GL, Tang YB, et al. Deficiency of volume-regulated ClC-3 chloride channel attenuates cerebrovascular remodelling in DOCA-salt hypertension. Cardiovasc Res. 2013;100(1):134–42. https://doi.org/10.1093/cvr/cvt156 PMID: 23786998

62. Wen J, Bao M, Tang M, He X, Yao X, Li L. Low magnitude vibration alleviates age-related bone loss by inhibiting cell senescence of osteogenic cells in naturally senescent rats. Aging (Albany NY). 2021;13(8):12031–45. https://doi.org/10.18632/aging.202907 PMID: 33888646

63. Vorhees CV, Williams MT. Morris water maze: procedures for assessing spatial and related forms of learning and memory. Nat Protoc. 2006;1(2):848–58. https://doi.org/10.1038/nprot.2006.116 PMID: 17406317

64. Wang F, Lu J, Peng X, Wang J, Liu X, Chen X, et al. Integrated analysis of microRNA regulatory network in nasopharyngeal carcinoma with deep sequencing. J Exp Clin Cancer Res. 2016;35:17. https://doi.org/10.1186/s13046-016-0292-4 PMID: 26795575

65. Lin C, Hsu Y, Huang Y, Shih Y, Wang C, Chiang W, et al. A KDM6A-KLF10 reinforcing feedback mechanism aggravates diabetic podocyte dysfunction. EMBO molecular medicine. 2019;11(5). https://doi.org/10.15252/emmm.201809828 PMID: 30948420

66. Qu Q, Wang L, Bing W, Bi Y, Zhang C, Jing X, et al. miRNA-126-3p carried by human umbilical cord mesenchymal stem cell enhances endothelial function through exosome-mediated mechanisms in vitro and attenuates vein graft neointimal formation in vivo. Stem Cell Res Ther. 2020;11(1):464. https://doi.org/10.1186/s13287-020-01978-z PMID: 33138861

67. Baciu C, Sage A, Zamel R, Shin J, Bai XH, Hough O, et al. Transcriptomic investigation reveals donor-specific gene signatures in human lung transplants. Eur Respir J. 2021;57(4). https://doi.org/10.1183/13993003.00327-2020 PMID: 33122335

68. Radavelli-Bagatini S, Blekkenhorst LC, Sim M, Prince RL, Bondonno NP, Bondonno CP, et al. Fruit and vegetable intake is inversely associated with perceived stress across the adult lifespan. Clin Nutr. 2021;40(5):2860–7. https://doi.org/10.1016/j.clnu.2021.03.043 PMID: 33940399

69. Rubega M, Formaggio E, Di Marco R, Bertuccelli M, Tortora S, Menegatti E, et al. Cortical correlates in upright dynamic and static balance in the elderly. Sci Rep. 2021;11(1):14132. https://doi.org/10.1038/s41598-021-93556-3 PMID: 34238987

70. Fedintsev A, Kashtanova D, Tkacheva O, Strazhesko I, Kudryavtseva A, Baranova A, et al. Markers of arterial health could serve as accurate non-invasive predictors of human biological and chronological age. Aging (Albany NY). 2017;9(4):1280–92. https://doi.org/10.18632/aging.101227 PMID: 28455973

